# Clathrin-mediated endocytosis of ERECTA family receptors is essential for proper stomatal development in Arabidopsis

**DOI:** 10.1101/2025.07.31.668009

**Authors:** Chi Zhang, Liang Chen, Suomin Wang, Chao Wang, Jianwei Pan, Suiwen Hou

## Abstract

Stomata, specialized structures of plant epidermis, are crucial for gas (e.g. CO_2_, O_2_ and H_2_O) exchange between plants and the environment. Stomatal density and pattern are governed by ERECTA Family (ERf) receptor-like kinases (RLKs)-controlled signaling. Clathrin-mediated endocytosis (CME) is a new mechanism for the internalization of receptor to activate the downstream signaling in animals. Here, we found that mutation of CME components resulted in formation of stomatal clusters and thus increased stomatal index. The adaptor protein 2 σ (AP2σ) subunit of CME interacted with ERf. ERf receptor internalization required CME. Furthermore, the ERf receptor motifs were identified to be recognized by AP2σ. Genetic analysis showed that CME components CHC2, CLC2, and CLC3 acted downstream of EPIDERMAL PATTERNING FACTOR 1/2 (EPF1/2) while upstream of YODA, and worked together with ERf to regulate stomatal development. Consistently, EPF2-induced MAPK activation was significantly reduced in *chc2-2 clc2 clc3* and *ap2μ clc2 clc3* mutants or when the ERf endocytic motifs were mutated, leading to stabilized SPEECHLESS protein and misregulated stomatal lineage progression. Overall, our findings demonstrate that activation of the ERf receptors via internalization is essential for stomatal development, establishing a novel mechanism of CME-mediated receptor internalization activation in plants.

**Significance Statement:** Stomatal development in Arabidopsis is well characterized, but how EPF-ERf signals activate the MAPK cascade remains unclear. While animals employ clathrin-mediated endocytosis (CME) for receptor internalization and signaling, this mechanism was unknown in plants. We identified CME as a novel regulator of stomatal development. The ERf receptors represent the first identified cargo proteins for AP2σ in plants, undergoing internalization through CME. Clathrin acts downstream of EPF but upstream of YODA, and blocking ERf internalization impairs MAPK activation. Thus, ERf internalization is essential for downstream signaling, revealing a novel CME-dependent activation mechanism in plants.

## Introduction

Stomatal development in Arabidopsis adheres to a “one-cell spacing rule”, wherein one or more pavement cells must separate two stomatal cells (1, 2). This spatial arrangement is crucial for optimizing the balance between CO_2_ uptake and water transpiration, thereby ensuring efficient stomatal function. Stomatal development progresses through three sequential transitions mediated by master bHLH transcription factors in Arabidopsis. SPEECHLESS (SPCH) initiates stomatal lineage commitment by driving asymmetric division of meristemoid mother cells (MMCs) into meristemoids (M) and stomatal lineage ground cells (SLGCs). MUTE terminates proliferative M divisions and triggers differentiation into guard mother cells (GMCs). FAMA executes the final transition by promoting GMCs symmetric division and guard cells (GCs) maturation (3, 4). These core regulators require heterodimerization with SCREAM (SCRM) to coordinate stomatal development (5–7).

The “one-cell spacing” pattern of Arabidopsis stomata is largely controlled by stomatal receptor-mediated signaling. The ERECTA Family (ERf) receptor-like kinases (RLKs), comprising ERECTA, ERECTA-LIKE 1 (ERL1), and ERL2, function as main stomatal receptors (8). ERECTA, ERL1, and ERL2 exhibit distinct roles in stomatal development. ERECTA and ERL2 function similarly, primarily acting during the early stage to suppress MMC division (9). In contrast, ERL1 specifically inhibits the differentiation of M into GMCs (10). ERf receptors recognized EPIDERMAL PATTERNING FACTOR (EPF), small secreted polypeptides that prevent or promote stomatal formation (11, 12). ERECTA can recognize EPF2, which activates downstream MITOGEN-ACTIVATED PROTEIN KINASE (MAPK) cascade represented by the phosphorylation of MPK3/6 (13–15). Then, MPK3/6 phosphorylate SPCH, ultimately leading to degradation of SPCH, and thus inhibiting the initiation of stomatal lineage (15, 16). On the other hand, ERL1-mediated recognition of EPF1 suppresses *MUTE* expression, and disruption of this signaling pathway leads to excessive MUTE accumulation that accelerates stomatal differentiation and causes aberrant stomatal patterning (11). ERECTA and ERL1 can also perceive EPF LIKE 9 (EPFL9/STOMAGEN), a mesophyll-derived antagonist of EPF1/2, but can’t activate MAPK cascade, resulting in the promotion of stomatal formation (17, 18). The ERf receptors require their co-receptor TOO MANY MOUTH (TMM) and SOMATIC EMBRYOGENESIS RECEPTOR-LIKE KINASES (SERKs) to properly regulate stomatal patterning and development (17, 19). Moreover, emerging studies have started to uncover the regulatory machinery governing ERECTA subcellular trafficking. Xue et al. identified an NRPM protein that promotes plasma membrane localization of ERECTA, enhancing its function (20). Loss of the ERDJ3B chaperone impairs ER quality control, reducing ERECTA protein abundance (21). PUB30/31-mediated ubiquitination targets ERECTA for vacuolar trafficking and degradation, attenuating its signaling (22, 23).

While stomatal patterning in Arabidopsis has been extensively characterized over 30 years, a fundamental question persists: how do ERf receptors relay EPF signals to the MAPK cascade? We break down this question into two sub-questions: How does ERf transmit the EPFs signal? And to whom/what downstream component does ERf relay the signal? Qi et al. established that ERL1 undergoes internalization, a process facilitated by externally applied ligands (24). It is well known that receptors, e.g., EPIDERMAL GROWTH FACTOR RECEPTOR (EGFR), PLATELET-DERIVED GROWTH FACTOR RECEPTOR (PDGFR), NEUROTENSIN RECEPTOR-1 (NTR1), are internalized through clathrin-mediated endocytosis (CME) to activate signaling pathways in animals (25, 26). Functioning as a highly conserved eukaryotic trafficking mechanism, CME drives membrane protein internalization via the dynamic assembly of clathrin-coated pits (CCPs) and their maturation into clathrin-coated vesicles (CCVs) (27). The process of CME requires the participation of many proteins, mainly including clathrin, adaptor proteins, and some accessory proteins (28). Clathrin function requires assembly of a cage-like complex composed of three clathrin heavy chains (CHCs) and three clathrin light chains (CLCs). *CHC* has two homologous genes *CHC1* and *CHC2*, and three *CLC* homologous genes *CLC1*, *CLC2* and *CLC3* in Arabidopsis (29–31). There are two main types of adaptor proteins: TPLATE complex (TPC) and adaptor protein 2 (AP2) complex (32, 33). TPC has eight subunits that mediate plasma membrane bending during CME, including TPLATE, TML etc. (33). AP2 is a heterotetrametric complex consisting of two large subunits (AP2α and AP2β), one medium subunit (AP2μ), and one small subunit (AP2σ) (32, 34). AP2μ and AP2σ recognize specific endocytic cargo motifs respectively, such as TyrXXΦ (YXXΦ, Φ represents a bulky hydrophobic amino acid) and Glu/AspxxxLeu/Ile (D/ExxxL/I) (35, 36). In plants, CME regulates the internalization of diverse cargo proteins, e.g., BRASSINOSTEROID INSENSITIVE 1 (BRI1), FLAGELLIN-SENSING 2 (FLS2), EF-TU RECEPTOR (EFR) and FERONIA (FER), thereby modulating a range of developmental and physiological processes (34, 37–48). However, it remains unclear whether ERf internalization occurs via CME and whether this process is essential for downstream signal transduction with regard to stomatal development.

Although CME functions broadly in plant growth, development, and stress responses, its role in stomatal development remains unknown. In this study, we identified “CME” as a novel stomatal regulatory factor. CME component mutants exhibit severe stomatal clustering, a phenotype partially resembling that of the *er105 erl1 erl2* mutants. We demonstrate that CME mediates ERf endocytosis and acts downstream of EPF1/2 but upstream of YODA in stomatal signaling. Consistently, CME impairment suppresses EPF2-induced MAPK activation, leading to SPCH stabilization and enhanced stomatal production. Our findings establish that CME is essential for downstream signaling activation and proper stomatal patterning, proposing a new mechanism of CME-mediated receptor internalization activation in plants.

## Results

### Clathrin and AP2 complex are involved in stomatal development

To study the role of CME in stomatal patterning, we systematically analyzed mutants of key CME components in Arabidopsis, including clathrin subunits (CHC2, CLC2, CLC3) and adaptor protein subunits (AP2μ, AP2σ). Our phenotypic characterization revealed that *ap2μ*, *ap2σ*, and *chc2-2* single mutants exhibited significant stomatal clustering and increased stomatal index compared to wild-type (Col-0) (Fig. 1 *A*-*D*, *M*), disrupting stomatal development pattern. We further constructed higher-order mutants and found that *ap2σ ap2μ*, *ap2μ clc2 clc3*, *ap2σ clc2 clc3*, and *chc2-2 clc2 clc3* displayed additive phenotypes with enhanced stomatal clustering and higher stomatal index (Fig. 1 *F*-*I*, *M* and *N*), though *clc2 clc3* double mutants showed single mutant phenotypes (Fig. 1 *E*). Complementation with respective genomic constructs (*pAP2μ::AP2μ-GFP*, *pAP2σ::AP2σ-GFP, pCHC2::CHC2-GFP*) fully rescued the corresponding single mutant phenotypes (Fig. 1 *J*-*L* and *M*). These observations suggest that clathrin and adaptor proteins, as key components of CME, coordinate to regulate stomatal development.

**Fig. 1.**
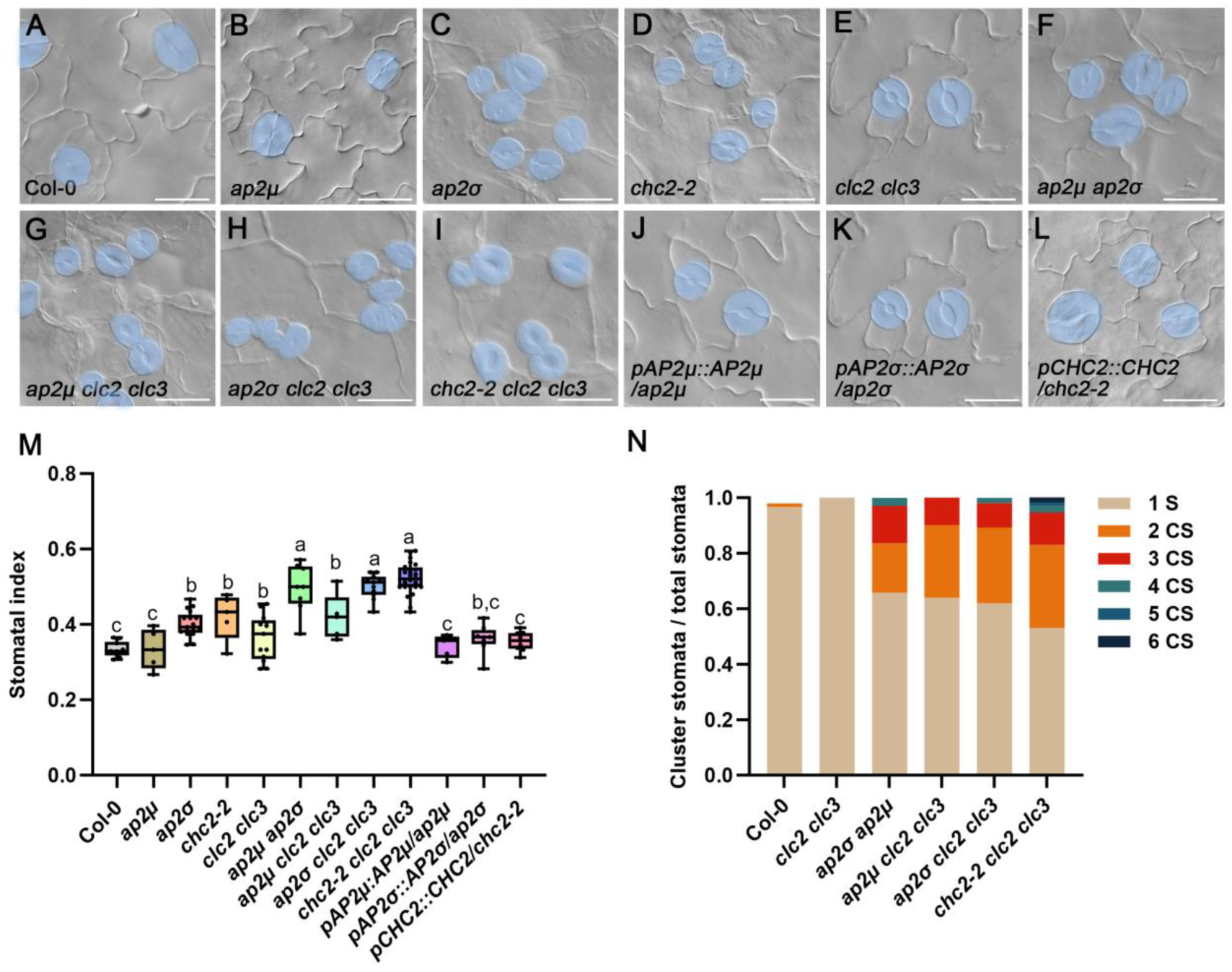
Arabidopsis clathrin and AP2 subunits mutations led to clustering stomata and increased stomatal index. (A-L) DIC images showed abaxial epidermal cell patterns on cotyledons at 5 days post-germination (dpg) in Col-0, *ap2μ*, *ap2σ*, *chc2-2*, *clc2 clc3*, *ap2μ clc2 clc3*, *ap2σ clc2 clc3*, *chc2-2 clc2 clc3*, *pAP2μ::AP2μ/ap2μ*, *pAP2σ::AP2σ/ap2σ* and *pCHC2::CHC2/chc2-2*. Mutations of clathrin and AP2 exhibited clustered stomata. The complementation plants of the single mutant resembled the wild type. Stomata were painted blue. Scale bars, 20 μm. (M) Statistical chart represented stomatal index in (A-L). Tukey test was determined for significance, values were mean ± SEM (n > 5). (N) Stacked bar graph showed the proportion of cluster stomata in (E-I). 1 S represent one stoma; 2-6 CS represent two-six clustered stomata.

To determine which specific stage of stomatal development is disrupted in CME component mutants, we analyzed the expression patterns of key stomatal lineage transcription factors ( *SPCH*, *MUTE*, *FAMA*) in *ap2μ*, *ap2σ*, *chc2-2*, *clc2 clc3*, and *chc2-2 clc2 clc3* mutants. We observed that ectopic *MUTE* expression in adjacent M cells frequently leading to paired GMCs expressing *FAMA* (Fig. S*1*), which explain the formation of stomatal clusters in CME mutants and suggest that CME primarily functions during early stomatal development to maintain proper lineage progression.

TPC, another adaptor protein, reaches the plasma membrane earlier than AP2, and then they colocalize there during early CME, suggesting they may cooperate in cargo protein recognition (33, 49, 50). To investigate the potential involvement of the TPC in CME-dependent stomatal development, we employed an estradiol (ESTR)-inducible downregulation system to bypass pollen lethality caused by TPC subunit mutations. As a result, neither *amiR-TPLATE* nor *amiR-TML* lines showed significant alterations in stomatal patterning or density compared to Col-0 (Fig. S*2*), suggesting that TPC is not essential for stomatal pattern in Arabidopsis.

### AP2σ subunit interacts with ERf receptors

Given CME’s role as a conserved eukaryotic pathway mediating selective membrane protein internalization, we examined its involvement in regulating stomatal receptors through trafficking. ERf receptors represent compelling candidates among stomatal signaling receptors, as evidenced by phenotypic similarity between the *chc2-2 clc2 clc3* and *er105 erl1 erl2* mutants (Fig. S*3*) and EPF1-induced ERL1 internalization (24). To determine if ERf receptors undergo CME-dependent trafficking, we initially examined their spatiotemporal association with the AP2 complex. AP2 σ, one subunit of the AP2 complex, was selected as representative. In tobacco epidermal cells, ERf receptors predominantly localized to the plasma membrane, while AP2σ exhibited diffuse cytoplasmic distribution with distinct plasma membrane accumulation (Fig. S*4*). Intriguingly, truncation analysis revealed that the cytoplasmic domain of ERf (ERf_CD_) localized to both the cytoplasm and nucleus yet maintained robust co-localization with AP2σ (Fig. S*4*). These results further imply that ERf receptors are likely recognized as cargoes by the AP2, subsequently internalized via the CME pathway.

In mammals, AP2σ and AP2μ have been characterized as core subunits responsible for cargo recognition in the adaptor protein complex (51). In plants, current evidence indicates that plants exclusively rely on AP2μ for cargo recognition. Therefore, we next employed a multi-platform approach to detect whether ERf could interact with AP2μ and AP2σ. Unexpectedly, yeast two-hybrid (Y2H) and bimolecular fluorescence complementation (BiFC) assays showed that ERf_CD_ receptors interact with AP2σ, but not AP2μ (Fig. 2 *A* and *B*). Co-immunoprecipitation (Co-IP) and *in vitro* pull-down assays further confirmed the AP2σ-ERf_CD_ interaction (Fig. 2 *C* and *D*). Collectively, these results demonstrate that the cytosolic domain of ERf receptors is directly recognized by AP-2σ, establishing ERf receptors as the first identified cargo proteins of AP2σ in plants.

**Fig. 2.**
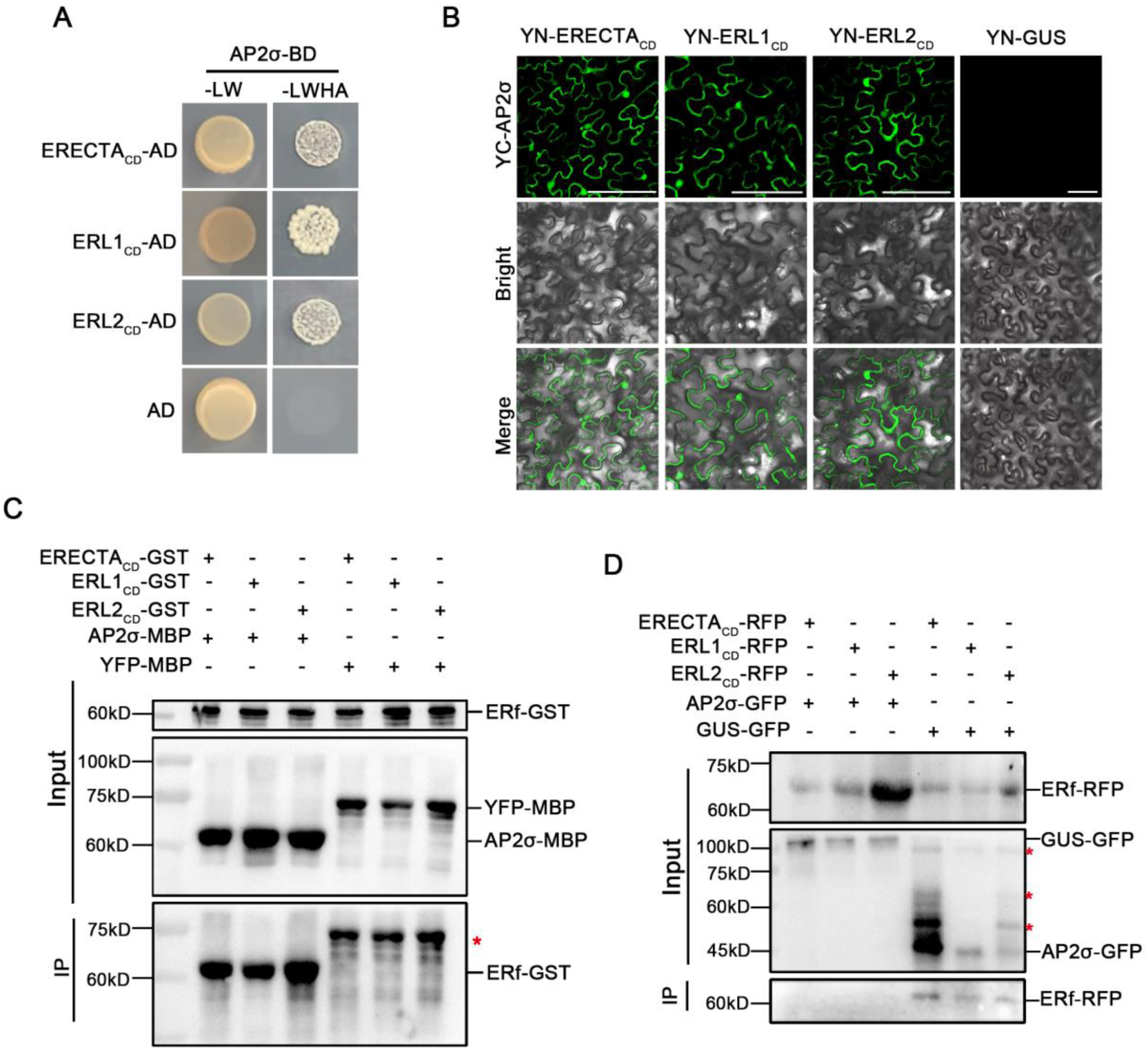
AP2σ subunit directly interacted with ERECTA_CD_, ERL1_CD_ and ERL2_CD_. (A) Y2H assayed the interaction of AP2σ with ERECTA_CD_, ERL1_CD_ and ERL2_CD_. “-LW” represents yeast two-deficiency medium (SD/-Trp/-Leu), “-LWHA” represents four-deficiency medium (SD/-Trp/-Leu/-His-Ade), and AD represents pGADT7 empty. (B) BiFC verified the interaction of AP2σ with ERECTA_CD_, ERL1_CD_ and ERL2_CD_, YN-GUS represents negative control. Scale bar, 100 μm. (C) Pull-down was used to verify the interaction between AP2σ and ERECTA_CD_, ERL1_CD_ and ERL2_CD_, and YFP-MBP was used as a negative control. Input represented the loading amount control. Red asterisks indicate nonspecific bands. (D) Co-IP was used to verify the interaction between AP2σ and ERECTA_CD_, ERL1_CD_ and ERL2_CD_, and GUS-GFP was used as a negative control.

### The endocytosis and subcellular localization of ERECTA/ERL1 are mediated by clathrin

To investigate whether ERf receptors undergo CME, we assessed the consequence of CME disruption on their internalization. FM4-64 is a highly lipophilic, water-soluble styryl dye that specifically binds to cell membranes to produce fluorescence, and widely used for labeling endocytic and exocytic membrane structure. FM4-64 labeling showed that wild-type stomatal precursor cells contained approximately 5 endosomal compartments per cell. In contrast, *chc2-2 clc2 clc3* mutant showed severe endocytic defects. Most cells completely lacked compartments, with only a minority forming 1-2 compartments (Fig. S*5*). These results establish clathrin’s essential role in endocytosis.

Next, we assessed whether the internalization of ERf receptors was affected in *chc2-2 clc2 clc3* mutants. Endocytosed cargo proteins are either recycled back to the plasma membrane for reuse or transported to the vacuole for degradation. Brefeldin A (BFA), a macrocyclic lactone antibiotic, inhibits protein trafficking by blocking secretory and membrane protein transport from the endoplasmic reticulum (ER) to the Golgi apparatus. The result showed that BFA treatment of *er105* plants expressing *ERECTA-YFP* induced reversible BFA body formation, whereas the *chc2-2 clc2 clc3* triple mutants displayed no detectable BFA bodies (Fig. 3 *A*). ERL1 showed deficient BFA body formation in *chc2-2 clc2 clc3*, phenocopying ERECTA’s trafficking defect (Fig. 3 *B*). These data indicate clathrin-dependent recycling is essential for plasma membrane retrieval of ERf receptors. Wortmannin (Wm), a specific inhibitor of phosphatidylinositol 3-kinase (PI3K), effectively blocks protein transport to the vacuole. While Wm treatment triggered robust responses in *er105* (44.3 ± 4.09% forming Wm bodies vs 0% baseline), *chc2-2 clc2 clc3* exhibited significantly attenuated accumulation (33.5 ± 6.1% vs 27.4 ± 4.7% baseline; p=0.0828, Student’s t-test) (Fig. 3 *C* and *D*). Similar to ERECTA, ERL1 showed impaired Wm body formation in the clathrin-deficient mutants (Fig. 3 *E* and *F*). These findings indicate that clathrin is also essential for proper ERf receptor internalization and subsequent sorting into PI3K-mediated degradative pathways.

**Fig. 3.**
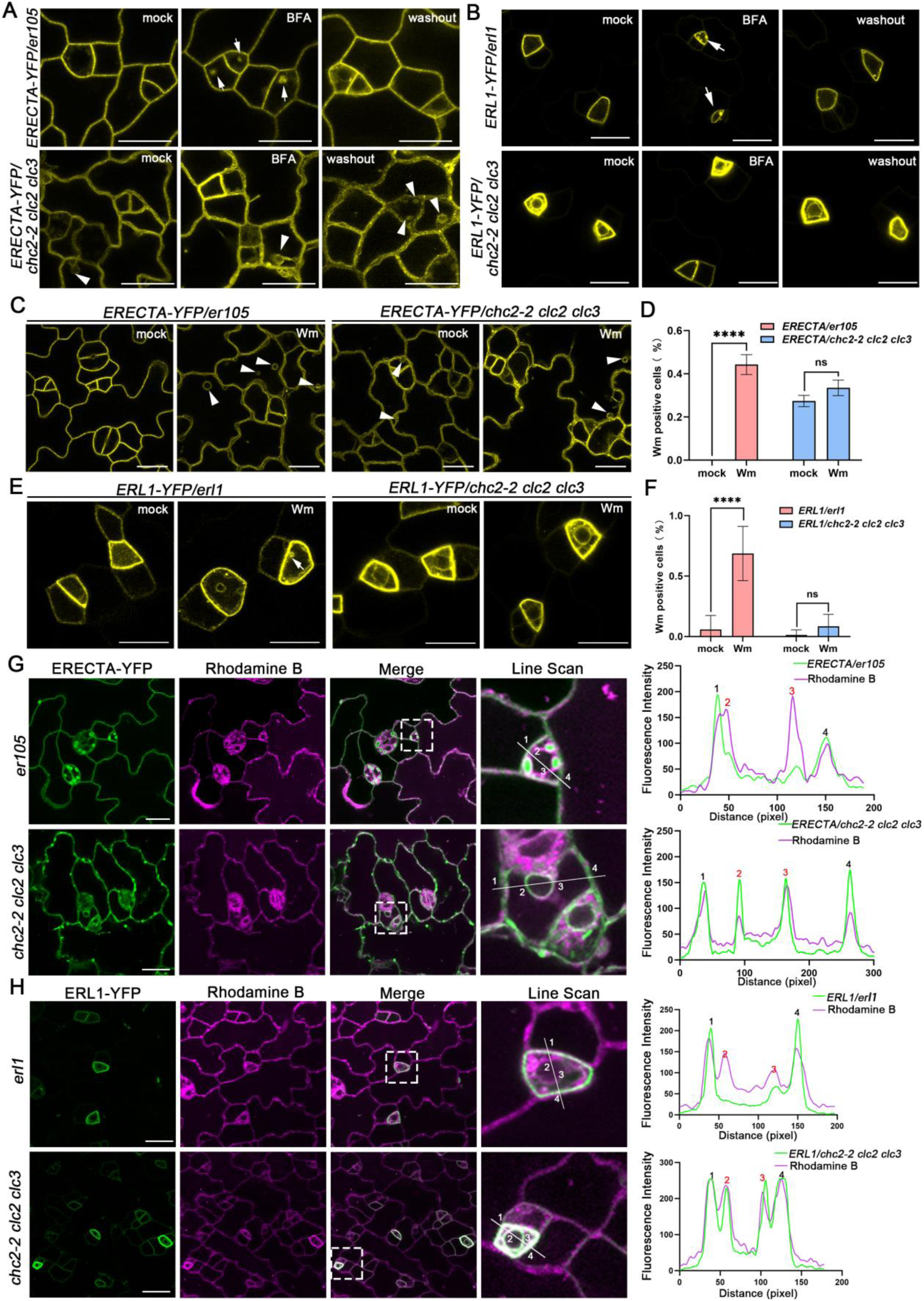
ERf receptors failed to undergo efficient internalization in *chc2-2 clc2 clc3* mutants, causing its accumulation in the plasma membrane and ER. (A) Confocal images showing the endocytosis of ERECTA in epiderma cells. *pERECTA::ERECTA-YFP* labeled *er105* and *chc2-2 clc2 clc3* seedlings were treated with 50 μM BFA for 2 h, followed by a 2 h washout with liquid 1/2 MS. Mock referring adding 0.4% DMSO, BFA representing 50 μM BFA for 2 h, white arrow indicating BFA bodies, washout representing 50 μM BFA for 2 h and washout BFA with liquid 1/2 MS for 2 h, white triangle pointing ring-like structures of ERECTA. Scale bars, 20 μm. (B) Confocal images showing the endocytosis of ERL1 in epiderma cells. *pERL1::ERL1-YFP* labeled *erl1* and *chc2-2 clc2 clc3* seedlings were treated with 50 μM BFA for 2 h, followed by a 2 h washout with liquid 1/2 MS. Scale bars, 20 μm. (C) Confocal images showing the vacuolar trafficking of endocytosed ERECTA. *pERECTA::ERECTA-YFP* labeled *er105* and *chc2-2 clc2 clc3* seedlings were treated with 50 μM Wm for 6 h. Images labeled “mock” show seedlings treated with 0.4% DMSO, while “Wm” images represent treatment with 50 μM Wm for 6h. Scale bars, 20 μm. (D) Statistical chart represented the proportion of cells containing Wm bodies among total cells in (C). Student’s *t*-test was determined for significance, values were mean ± SEM (n > 10). **** indicated *P*<0.0001, ns indicated no signification difference. (E) Confocal images showing the vacuolar trafficking of endocytosed ERL1. *pERL1::ERL1-YFP* labeled *erl1* and *chc2-2 clc2 clc3* seedlings were treated with 50 μM Wm for 10 min. Images labeled “mock” show seedlings treated with 0.4% DMSO, while “Wm” images represent treatment with 50 μM Wm for 10 min. Scale bars, 20 μm. (F) Statistical chart represented the proportion of cells containing Wm bodies among total cells in (E). Student’s *t*-test was determined for significance, values were mean ± SEM (n > 10). **** indicated *P*<0.0001, ns indicated no signification difference. (G) Confocal images showed that colocalization of ERECTA-YFP with the ER dye Rodamine B in *chc2-2 clc2 clc3* stomatal precursor cells. And colocalization was confirmed by quantitative analysis of fluorescence intensity profiles. Line scan is an enlarged image of the white dotted line box in the merge diagram. 1 and 4 representing plasma membrane, 2 and 3 represent ER. Scale bars, 20 μm. (H) Confocal microscopy revealed colocalization of ERL1-YFP with the ER dye Rhodamine B in *chc2-2 clc2 clc3* stomatal precursor cells. This colocalization was confirmed by quantitative analysis of fluorescence intensity profiles. The line scan image is an enlarged view of the region indicated by the white dotted box in the merge panel. 1 and 4 representing plasma membrane, 2 and 3 representing ER. Scale bars, 20 μm.

Surprisingly, clathrin-deficient mutants exhibited pronounced accumulation of ERECTA and ERL1 on the plasma membrane and formed ring-like structures in the cytoplasm (Fig. 3 *A*-*D*). These ring-like structures showed precise colocalization with the ER marker dye Rhodamine B (Fig. 3 *G* and *H*), indicating ERf receptor mislocalization to the ER in *chc2-2 clc2 clc3*. These phenotypes were faithfully recapitulated by the CME inhibitor ES9-17 (Fig. S*7*), demonstrating that CME activity is required to prevent ER retention of ERf receptors. These observations indicate that CME disruption impedes ERf receptor internalization from the plasma membrane or transport from ER to plasma membrane.

### The endocytosis motifs of ERf receptors govern their endocytic trafficking and subcellular localization

Identification of the AP2σ and ERf receptor interaction (Fig. 2) prompted us to investigate whether plants employ conserved D/ExxxL/I endocytic motifs for AP2 recruitment, as observed in mammals (27). Bioinformatics analysis predicted five putative AP2σ binding motifs in the cytoplasmic domain of ERf receptors, all featuring the canonical D/ExxxL/Ι sequence (Fig. S*8*). To test their functional relevance, we generated mutated version of ERf_CD_ (ERf_CD_-m) where all five motifs were replaced by AlaxxxAla (AxxxA) substitutions. Y2H and BiFC assays showed complete abolition of AP2σ interacting with ERf_CD_-m (Fig. 4 *A* and *B*), demonstrating that these motifs are indispensable for AP2σ recognition in planta.

**Fig. 4.**
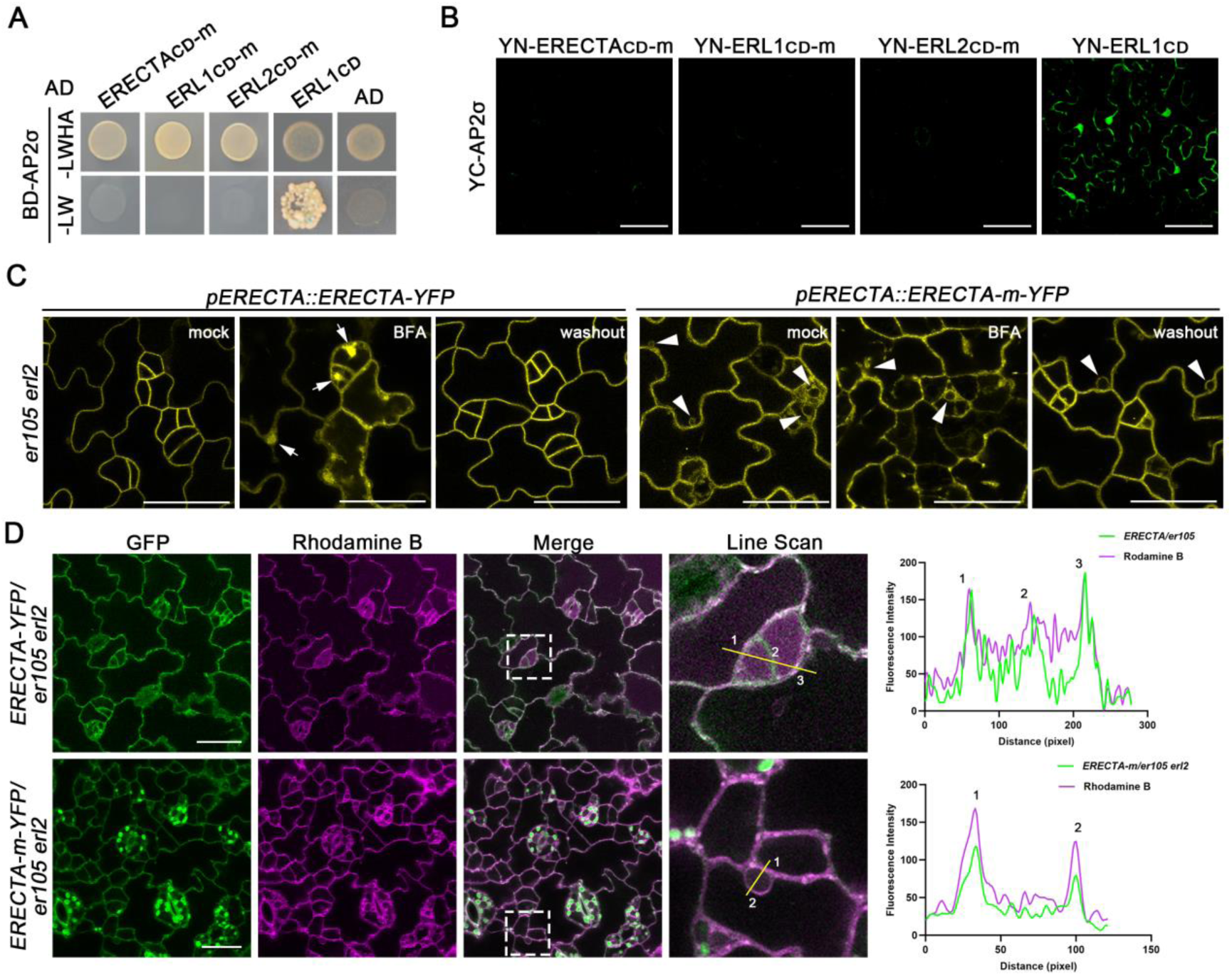
The mutation in ERf’s receptors AP2σ-binding motif disrupted its interaction with AP2σ, impairing endocytosis and causing ER retention. (A) Y2H assay showed the interaction of ERECTA_CD_-m, ERL1_CD_-m and ERL2_CD_-m with AP2σ. Unmutated ERL1_CD_ is a positive control. “-LW” represents yeast two-deficiency medium (SD/-Trp/-Leu), “-LWHA” represents four-deficiency medium (SD/-Trp/-Leu/-His-Ade), and AD represents pGADT7 empty. (B) BiFC assay showed the interaction between AP2σ and ERECTA_CD_-m, ERL1_CD_-m and ERL2_CD_-m in tobacco epidermis. Unmutated ERL1_CD_ is a positive control. Scale bars, 100 μm. (C) Confocal images showing the endocytosis of ERECTA in epiderma cells. *pERECTA::ERECTA-YFP* and *pERECTA::ERECTA-m-YFP* in *er105 erl2* mutants were treated with 50 μM BFA for 2h, followed by a 2-hour washout with liquid 1/2 MS. Mock referring adding 0.4% DMSO. BFA representing 50 μM BFA for 2h, white arrow indicates the BFA body, washout representing 50 μM BFA for 2 h and washout BFA with liquid 1/2 MS for 2 h, white triangle pointing ring-like structures of ERECTA. Scale bars, 20 μm. (D) Confocal images showed that colocalization of ERECTA-YFP with the ER dye Rodanmine B in *pERECTA::ERECTA-m-YFP/er105 erl2* stomatal precursor cells. And colocalization was confirmed by quantitative analysis of fluorescence intensity profiles. Line scan is an enlarged image of the white dotted line box in the merge diagram. 1 represented plasma membrane, and 2 was ER. Scale bars, 20 μm.

Next, to investigate whether the D/ExxxL/I motif mediates ERf receptor endocytosis, we generated an *ERECTA-m* plant (*pERECTA::ERECTA-m-YFP*) in which all D/ExxxL/I motifs were changed to AxxxA sequences in the genomic locus and introduced it into the *er105 erl2* double mutants. Stomatal observation in *ERECTA-m* showed severe clustering of stomatal precursor cells, with no phenotypic recovery of *er105 erl2*, resembling the phenotype of *epf2* mutant (Fig. S*9*).

These results indicate that mutation in the endocytic motif of ERECTA impairs its function. Subsequently, we examined whether the endocytic activity and subcellular localization of *ERECTA-m* were altered. After BFA treatment, BFA body formation was completely absent in *ERECTA-m* plants (Fig. 4 *C*). Moreover, ring-like structures, mirroring that of ERECTA observed in *chc2-2 clc2 clc3* mutant (Fig. 4 *D*), were observed in *ERECTA-m* plants. These results demonstrate that mutation of the AP2σ-binding motifs, responsible for mediating AP2σ’s specific recognition of ERf receptors, results in blocked endocytosis and retention of ERECTA in the ER.

### Genetic analysis of clathrin in the stomatal developmental pathway

To elucidate the genetic relationship between clathrin and ERf signaling in stomatal development, we generated higher-order mutants by crossing *chc2-2 clc2 clc3* with *er105 erl1 erl2*. Compared with the *er105* single mutant, the *chc2-2 clc2 clc3 er105* quadruple mutants only exhibited a slight decrease in the proportion of immature stomatal lineage cells (Fig. 5 *A*, *B*, *E*, *F* and *I*). In contrast to the *er105 erl2* double mutants, the *chc2-2 clc2 clc3 er105 erl2* quintuple mutants showed higher stomatal density but significantly reduced proportion of immature stomatal lineage cells (Fig. 5 *C*, *G*, *I* and *J*). However, no significant differences were observed in stomatal density or the proportion of immature stomata between the *chc2-2 clc2 clc3 er105 erl1 erl2* sextuple mutants and the *er105 erl1 erl2* triple mutants (Fig. 5 *D*, *H*, *I* and *J*). These results indicate that clathrin likely functions synergistically with ERf receptors to facilitate stomatal development, which is consistent with the result that ERf receptors associate with the CME complex.

**Fig. 5.**
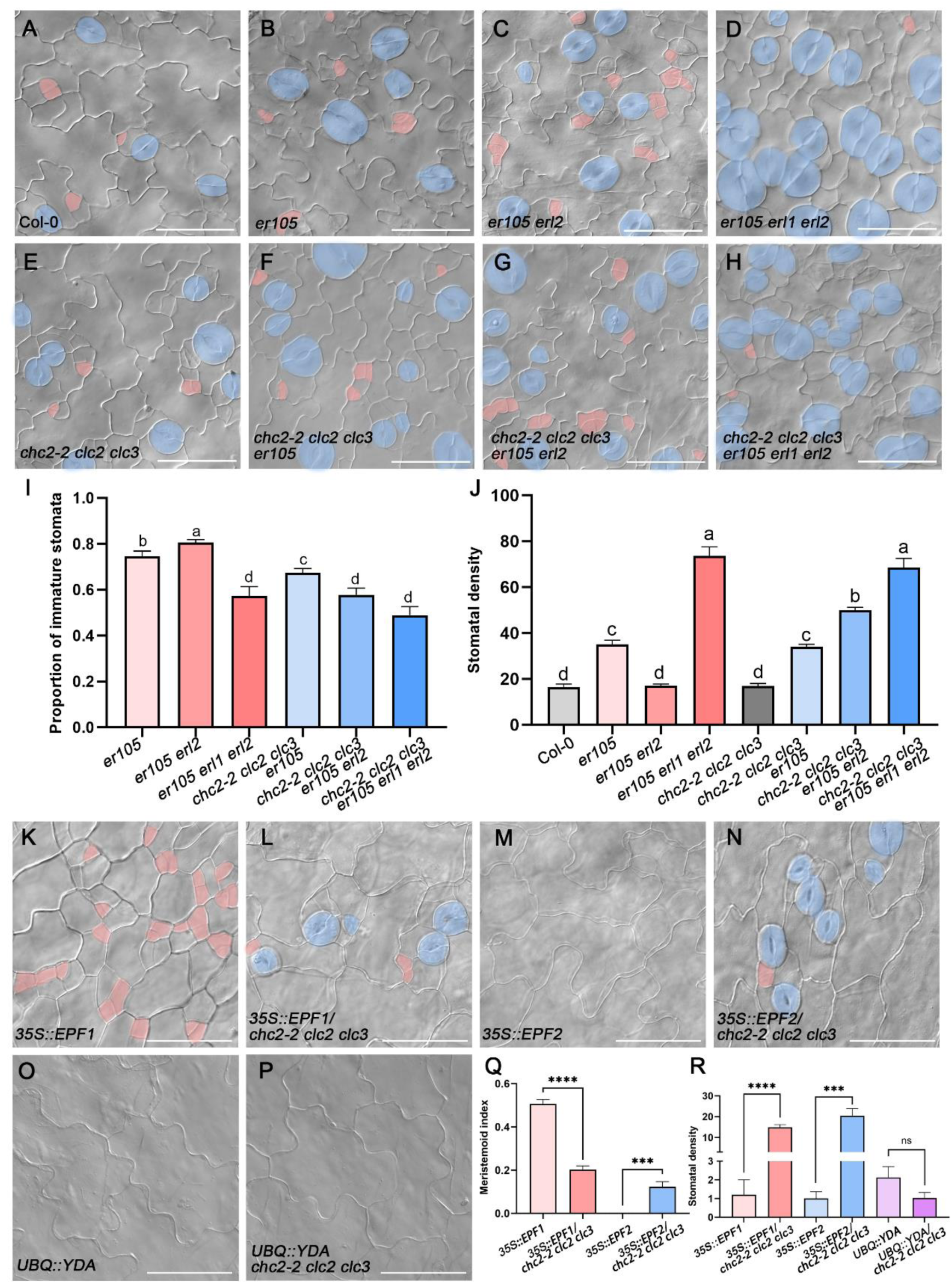
Clathrin was genetically located downstream of EPF1/2 and upstream of YODA, and formed a complex with ERf receptors to regulate stomatal development. (A-H) DIC images showed that abaxial epidermal cell patterns on cotyledons at 5 dpg in WT (top left column), *er105* (top second column), *er105 erl2* (top third column), *er105 erl1 erl2* (top right column), *chc2-2 clc2 clc3* (bottom left column), *chc2-2 clc2 clc3 er105* (bottom second column), *chc2-2 clc2 clc3 er105 erl2* (bottom third column) and *chc2-2 clc2 clc3 er105 erl1 erl2* (bottom right column). clathrin and ERf may act synergistically. For easy observation, the MMC or M is painted pink, and the stomatal are painted blue. Scale bars, 50 μm. (I) Statistical chart showed that the proportion of immature stomata was counted in B-D and F-H. Immature stomatal ratio = (stomatal lineage cell-stomatal cell)/stomatal lineage cell; (SLC-S)/SLC. Tukey test is determined for significance, values are mean ± SEM (n > 5). (J) Statistical chart showed that stomatal density of (A-H), statistical area is 0.1 mm^2^. Tukey test is determined for significance, values are mean ± SEM (n > 5). (K-P) DIC image showed that abaxial epidermal cell patterns on cotyledons at 4 dpg in *35S::EPF1* (top left column), *35S::EPF1/chc2-2 clc2 clc3* (top middle column), *35S::EPF2* (top right column), *35S::EPF2/chc2-2 clc2 clc3* (bottom left column), *UBQ::YDA* (bottom middle column) and *UBQ::YDA/chc2-2 clc2 clc3* (bottom right column) seedling. Clathrin functioned downstream of EPF1 and EPF2 while being upstream of YODA. For easy observation, the MMC or M is painted pink, and the stomatal are painted bule. Scale bars, 50 μm. (Q) Quantitative analysis showed the M index of (K-N). Student’s *t*-test was determined for significance, values were mean ± SEM (n > 5). *** indicates *P*<0.001, **** indicates *P*<0.0001. (R) Statistical chart showed that stomatal density in (K-P), the area in the picture is 0.1 mm^2^. Student’s *t*-test is determined for significance, values are mean ± SEM (n > 5). *** indicates *P*<0.001, **** indicates *P*<0.0001, ns indicates P>0.05.

We further investigated the genetic interactions between clathrin and EPF1/2, as well as YODA in stomatal development. Overexpression *EPF1* in *chc2-2 clc2 clc3* did not induce the characteristic stomatal suppression phenotype observed in wild type (Fig. 5 *K*, *L*, *Q* and *R*). Similarly, *EPF2* overexpression in the mutants recapitulated the original mutant phenotype and did not induce the stomatal suppression (Fig. 5 *E*, *M*, *N*, *Q* and *R*). These results showed that clathrin functions downstream of EPF1 and EPF2. Consistent with effects in wild-type plants, overexpression *YODA* in *chc2-2 clc2 clc3* resulted in complete suppression of stomatal lineage progression and exclusive pavement cell production (Fig. 5 *E*, *O*, *P*, *Q* and *R*), indicating that clathrin acts upstream of YODA. Collectively, these findings establish that functional CME is essential for the proper EPF-ERf-YODA signaling during stomatal development.

### CME-controlled ERf receptor endocytosis is essential for activation of EPF signaling

To investigate how ERf receptor internalization contributes to EPF signaling, we analyzed endocytic trafficking of ERf receptors in *epf* ligand mutants that cannot bind the receptor and thus cannot initiate downstream signaling cascades. The results showed that whereas BFA treatment induced ERECTA-containing BFA bodies in wild type, it failed to do so in the *epf2* mutant (Fig. 6 *A*). Similarly, ERL1-containing BFA bodies were not observed in the *epf1* mutant following BFA treatment (Fig. 6 *B*), demonstrating that EPF-ERf receptor complex formation is prerequisite for the initiation of CME of the ERf receptors.

**Fig. 6.**
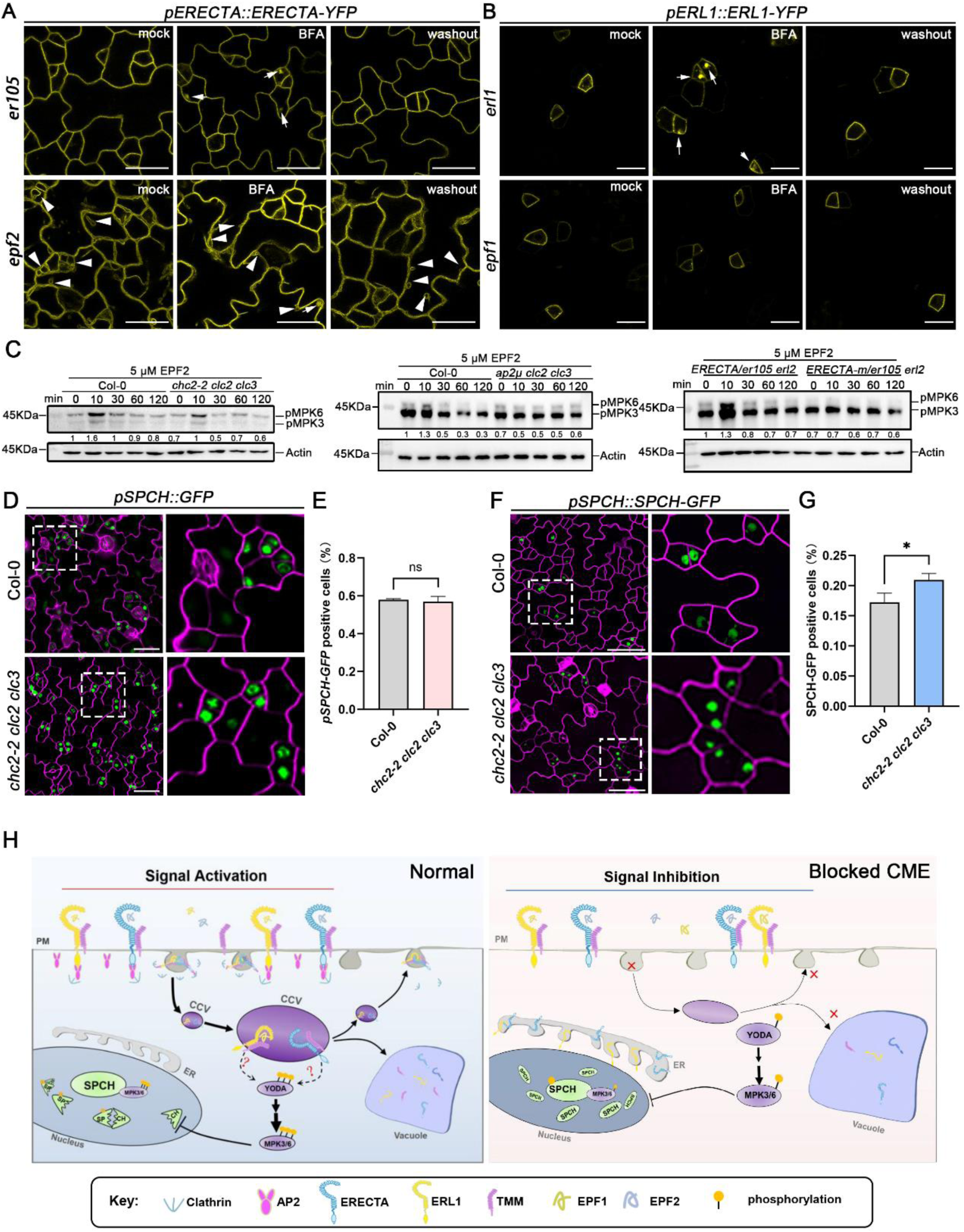
Impaired ERf receptor endocytosis compromised EPF signaling transmission. (A) Confocal images showing the endocytosis of ERECTA in epiderma cells. BFA represents the image after 50 μM BFA for 2 h, white arrow indicates the BFA body. Washout represents the image after 50 μM BFA for 2h and washout BFA with liquid 1/2 MS for 2 h. Scale bars, 20 μm. (B) Confocal images showing the endocytosis of ERL1 in epiderma cells. *pERL1::ERL1-YFP/erl1* and *pERL1::ERL1-YFP/epf1* seedlings were treated with 50 μM BFA for 1h, followed by a 1-hour washout with liquid 1/2 MS. Scale bars, 20 μm. (C) EPF2-triggered phosphorylation of MPK3/6 was diminished in *chc2-2 clc2 clc3*, *ap2μ clc2 clc3* and *ERECTA-m* plants. Arabidopsis seedlings of wild type and *chc2-2 clc2 clc3*, *ap2μ clc2 clc3* and *ERECTA-m* plants were grown under normal conditions for 12 days, then treated with 5 μM EPF2 for 0, 10, 30, 60, and 120 minutes. Phosphorylation levels of MPK3/6 (pMPK3/6) were analyzed by western blotting, with Actin serving as the loading control. (D) Confocal images showed that the expression of *pSPCH::GFP* in Col-0 and *chc2-2 clc2 clc3* mutants. FM4-64 was marked plasma membrane (magenta). The picture on the right is an enlarged picture of the white dotted box. Scale bar, 20 μm. (E) Quantitative results showed that the percentage of *pSPCH::GFP* positive cells. Data are represented as means ± SD (n>90 stomata from three independent experiments). Statistical significance was determined by Student’s *t*-test (ns, not significant, *p*>0.05). (F) Confocal images showed that the expression of *pSPCH::SPCH-GFP* in Col-0 and *chc2-2 clc2 clc3* mutants. FM4-64 was plasma membrane (magenta). The picture on the right is an enlarged picture of the white dotted box. Scale bar, 20 μm. (G) Quantitative analysis revealed the percentage of SPCH-GFP-positive cells. Data represent mean ± SD (n>90 stomata from three independent experiments). Statistical significance was determined by Student’s *t*-test (*p*<0.05; ns, *p*>0.05. (H) Schematic model of CME-mediated ERf receptors endocytosis regulating stomatal patterning. Under normal conditions, CME-localized in the cytoplasm can endocytose ERf, facilitating the transmission of upstream EPF-ERf signaling downstream to MAPK, thereby regulating the activity of SPCH. When CME is inhibited, the endocytosis of ERf is obstructed, preventing effective transmission of the EPF-ERf signal. Furthermore, the subcellular dynamics homeostasis of ERf is disrupted, leading to its retention in the ER and ultimately hindering the degradation of SPCH, resulting in the clustering stomata.

Since MPK3/6 phosphorylation status serves as both an indicator of upstream EPF-MAPK signaling activity and a predictor of downstream SPCH stability (15), we examined MPK3/6 phosphorylation dynamics in *chc2-2 clc2 clc3*, *ap2μ clc2 clc3* and *ERECTA-m* mutants following EPF2 treatment for signal initiation. Time-course analysis of MPK3/6 activation revealed that 5 μM EPF2 triggered transient phosphorylation in wild type, peaking at 10 min post-treatment (Fig. 6 *C*). In clathrin component deficient mutant *chc2-2 clc2 clc3*, MPK3/6 phosphorylation was significantly reduced despite maintaining wild-type-like activation kinetics (Fig. 6 *C*). By contrast, triple mutants lacking adaptor protein AP2μ together with clathrin components CLC2 and CLC3 exhibited almost abolition of both amplitude and activation kinetics of MPK3/6 (Fig. 6 *C*). Moreover, mutagenesis of the endocytic motif of ERECTA receptor completely eliminated MPK3/6 activation amplitude and temporal dynamics (Fig. 6 *C*). These results indicate that CME of ERECTA endocytosis is fundamentally required for EPF2-mediated activation of the MAPK cascades.

Further analysis of SPCH protein revealed a significantly higher proportion of SPCH-GFP-expressing cells in *chc2-2 clc2 clc3* mutants compared to wild type (Fig. 6 *F* and *G*). However, SPCH promoter activity showed no significant difference between the wild type and mutant (Fig. 6 *D* and *E*). These findings demonstrate that the reduced MPK3/6 phosphorylation levels, caused by impaired ERECTA endocytosis, enhance SPCH protein stability, ultimately leading to disordered stomatal patterning.

## Discussion

This work establishes CME as an essential regulator controlling stomatal development (Fig. 6 *H*). We demonstrate that the AP2σ subunit of CME recognizes ERf receptors, and clathrin disruption attenuates internalization of ERf receptors. Genetic analyses further reveal that clathrin acts downstream of EPF ligands but upstream of the YODA kinase in stomatal development signal pathway. Consistently, disruption of CME components or mutagenesis of the ERECTA endocytic motif blocks MPK3/6 activation. Collectively, these findings indicate that CME serves as a central hub that coordinates ERf endocytosis to activate MAPK signaling, thereby filling the gap between EPF-ERf receptors and MAPK cascade during signal transduction of stomatal development.

### CME is a novel regulator for stomatal development

CME serves as the primary mechanism for internalizing plasma membrane proteins. While investigating whether endocytosis of ERf receptors is mediated by CME, we unexpectedly observed that mutants of core CME components displayed a clustered stomata phenotype. And this phenotype shows similarities to that reported for *er105 erl1 erl2* mutants (10). We found that ERf receptor internalization was impaired in clathrin mutants, despite the fact that this process had been previously documented (9, 21, 24). This establishes that ERf receptor internalization is controlled by CME, a mechanism whose involvement was previously unclear. Clathrin requires collaboration with the adaptor protein AP2 complex to regulate stomatal development, though it may function independently in the nucleus to regulate DNA damage repair (52). Specifically, single mutants of clathrin heavy chain (*chc2-2*), as well as AP2 subunits (*ap2σ*, *ap2μ*), display stomatal clustering. Furthermore, double and triple mutants (*ap2σ chc2-2*, *ap2σ clc2 clc3*, *ap2μ clc2 clc3*, *chc2-2 clc2 clc3*) exhibit significantly more pronounced stomatal clustering. This progressive severity indicates that the extent of stomatal clustering is negatively correlated with residual CME activity.

In our study of ERf receptor endocytosis in the *chc2-2 clc2 clc3* and *ERECTA-m* mutants, we observed not only significantly inhibited internalization of ERf, but also accumulation of ERf receptors at the plasma membrane and unexpected retention in the ER. We propose two potential mechanisms for this aberrant ER localization. Firstly, disruption of CME may activate cellular quality control systems, causing ER retention: a phenomenon analogous to that mediated by *ERDJ3B* (21). Secondly, CME may be functionally coupled to clathrin-dependent secretory pathways (53), and its disruption could impair ERf receptor transport to the plasma membrane, resulting in ER accumulation.

### ERf is the first cargo protein recognized by AP2σ in plants

Adaptor proteins initiate CME by recognizing cargo proteins. In animals, only one class of AP2 was identified (28). However, plants harbor an additional conserved adaptor complex, TPC, alongside AP2 (33). The functional relationship between AP2 and TPC remains unclear. Structural and biochemical studies predict that TPC interacts with clathrin, AP2α, AP2σ, dynamin-related proteins (DRPs), and mediates membrane curvature during CME (33, 49, 50). Crystallographic analyses of AP2 reveal that cargo binding induces substantial conformational changes, which reposition its four phosphatidylinositol 4,5-bisphosphate (PtdIns(4,5)P₂) binding sites and two endocytic motif-binding sites into a coplanar configuration, thereby facilitating simultaneous interaction with both PtdIns(4,5)P₂ and cargo proteins (32). Unlike AP2, which regulates diverse biological processes through recognition of multiple cargo proteins, TPC is currently the only factor known to function in Arabidopsis shoot apical meristem maintenance by attenuating CLAVATA1 signaling (54). Our study demonstrates that the adaptor protein AP2 complex plays a predominant role in regulating stomatal development, whereas the TPC complex is largely dispensable. This is evidenced by the observation that mutations in AP2 subunits cause stomatal clustering, whereas functional impairment of TPC subunits does not result in discernible stomatal developmental defects.

Surprisingly, unlike cargo proteins in other plants that are recognized by the AP2 μ subunit, the ERf receptors are recognized by the AP2σ subunit to initiate CME. We confirmed the interaction between AP2σ and ERf receptors. More importantly, mutation of the specific motif within the cytoplasmic domain of ERf that is recognized by AP2σ abolished both the AP2σ-ERf interaction and ERf internalization. The ERf receptors are the first cargo protein in plants demonstrated to be recognized by AP2σ. This finding not only highlights the specificity of ERf recognition but also suggests functional conservation similar to mechanisms observed in animal cells. Further, in animals, phosphorylation of the AP2 subunit by kinases BMP-2-INDUCIBLE KINASE (BMP2K) and ADAPTOR-ASSOCIATED KINASE1 (AAK1) enhances its functionality (55, 56). Homolog of *AAK1* also exists in plants, and *aak1* mutant phenocopies *ap2μ* defects (44). Whether AP2σ is also regulated by protein phosphorylation is interesting to pursue in future.

### CME is necessary for signal transduction of ERf receptors, as well as some important RLKs in plants

This study establishes that ligand (EPF) binding triggers CME of the ERf receptor, enhancing EPF-ERf signaling through the MAPK cascade. Genetically, clathrin mutations disrupt EPF signal transduction and block YODA activation, demonstrating the essential role of clathrin in EPF signaling. Biochemically, EPF2-induced phosphorylation of MPK3/6 is significantly reduced in *chc2-2 clc2 clc3* mutants, confirming clathrin is required for downstream signaling. This is further validated by impaired MPK3/6 activation in *ap2μ clc2 clc3* mutants. Consistently, ERECTA endocytic motif mutations completely abolish MPK3/6 activation, supporting that CME is indispensable for EPF signal transmission. The differential phosphorylation of MPK3/6 likely arises from compromised functional integrity of CME. Because *ERECTA-m/er105 erl2* mutants exhibit complete blockade of ERECTA internalization, while *ap2μ clc2 clc3* mutants likely convey greater impairment of ERECTA endocytosis than *chc2-2 clc2 clc3*. Additionally, in *epf* mutants, ERf receptor internalization is impaired in stomatal lineage cells. This indicates that endocytosis of ERf receptors requires EPF ligand binding to transduce signals, and thus cannot occur in the absence of EPF. Therefore, ERf receptor trafficking is essential for their function in specifying stomatal cell fate.

In plants, the detailed mechanisms by which some important RLKs, such as FER, FLS2, RGF1 INSENSITIVES (RGI), etc., transmit signals into cytoplasmic and subsequently activate the MAPK cascade signal have not yet been completely resolved. Based on our data, we propose that CME-mediated receptor internalization activation not only plays an indispensable role in ERf receptor-regulated stomatal development, but also serves a similarly suitable function in diverse developmental processes and stress responses controlled by other RLKs. For instance, the short-root phenotype of *rgi* mutant resembles that of *ap2μ* mutant defective in CME components, suggesting that RGI receptor likely undergoes CME-dependent internalization to activate MAPK downstream signaling for root meristem development (44, 57). CME also regulates FER receptor internalization, with developmental defects and impaired stress responses in *fer* mutant partially phenocopying those observed in CME component mutants (58, 59). Likewise, CME regulates pattern-triggered immunity (PTI)-mediated by receptors including FLS2, EFR, and PLANT ELICITOR PEPTIDE RECEPTOR 1 (PEPR1) (40, 60, 61). Collectively, these findings also strongly hint at receptor internalization-dependent activation by CME as a fundamental yet underappreciated mechanism initiating downstream signaling in plants. Our work redefines CME role in signal transduction by elucidating how ligand-receptor complexes activate the MAPK cascade.

Although we have confirmed that EPF-ERf signaling to MAPK occurs through CME-dependent receptor internalization, it remains still unclear both at which stage of CME-internalized ERf receptors transmits signals and how this transduction directly or indirectly initiates MAPK cascade activation. We think that internalized ERf receptors trigger downstream signals within CCPs/CCVs in plants, similar to the mechanism in animals (25). Future work should pay attention to the detail mechanism and also how the endocytosed ERf receptors couples MAPK cascade.

## Acknowledgments

We are grateful to Dr. Juan Dong (University of Texas at Austin) for generously providing the seeds of ERECTA and ERL1 transgenic lines (*ERECTA-YFP; er105* and *ERL1-YFP; erl1*). We also sincerely thank Dr. Jijie Chai (Tsinghua University) for kindly sharing the EPF1 and EPF2 peptides. Additionally, we acknowledge the Arabidopsis Biological Resource Center (ABRC) for supplying the T-DNA insertion mutants. This research was supported by the following funding: The National Key Research and Development Program of China (Grant No. 2022YFD1201801), the National Natural Science Foundation of China (Grant No. 32170340), the Science and Technology Program of Gansu Province (Grants No. 22ZD6NA049, 21ZD10NF003 −2, 24JRRA428 and 24JRRA441), the Major Science and Technology Project of Gansu Province (Grant No. 17ZD2NA016), the Top Leading Talents Program of Gansu Province, and the Chang Jiang Scholars Program of China (2023).

## Author Contributions

C.Z. and S.W.H. conceived and designed the research project. C.Z. performed genetic analyses, immunoblotting experiments, and cell biological studies. The manuscript was written by C.Z., C.L., and S.W.H. All authors participated in manuscript editing and approved the final version. J.W.P. and S.W.H. provided funding and resources for this study.

## Materials and Methods

### Plant Material and Growth Conditions

The wild-type control used in this paper is Columbia (Col-0). Mutants *ap2μ* (SALK_083693c), *ap2σ* (SALK_141555), *chc2-2* (SALK_042321), *clc2* (SALK_016049), *clc3* (CS100219), *pAP2μ::AP2μ-GFP/ap2μ*, *pAP2σ::AP2σ-GFP*/*ap2σ* were provided by professor Jianwei Pan. *er105 erl1 erl2* is a gift from Professor Keiko U. Torii ’s; *pERECTA::ERECTA-YFP/er105* and *pERL1::ERL1-YFP/erl1* were donated by Dr. Juan Dong. The *35S::EPF1*, *35S::EPF2*, *UBQ::YDA*, *pSPCH::nucGFP*, *pMUTE::nucGFP*, *pFAMA::nucGFP*, *pSPCH::SPCH-GFP* were constructed in our laboratory. Sterilized seeds were grown on half strength of Murashige and Skoog (1/2 MS) medium supplemented with 1% sucrose and 1% agar at 22°C in a growth room with a 16 h light/8 h dark light regime.

### Plasmid Construction

Most plasmids used in this study were generated using a Gateway Cloning Kit (Thermo Fisher Scientific) according to the manufacturer’s instructions. Briefly, full-length coding sequence fragments of ERECTA, ERL1, ERL2, ERECAT_CD_, ERL1_CD_, ERL2_CD_, YODA, EPF1, EPF2 or SPCH and full-length genomic sequence ERECTA, ERL1, ERL2 were amplified and inserted into *pDONR-Zeo* via the Gateway BP Clonase I reaction. Sequence-confirmed coding sequences were then subcloned into destination vectors. The primer sequences are provided in Supplemental Data Tab. S1.

### DIC Imaging

Transparency: according to the ratio of anhydrous ethanol : acetic acid = 3 : 1 configuration of the required dose of transparent liquid. The 5-day-old Arabidopsis seedlings were transparent overnight in a 12-well plate; Alkalization: Absorb the transparent liquid after overnight transparency, add the configured alkalization solution (water : ethanol = 2 : 3, 7% NaOH ), alkalize for at least 4 h; Hydration: After sufficient alkalization, the alkalization liquid is sucked out. Hydrate with 40 %, 20%, 10% ethanol gradient respectively. Each stage was performed for at least 20 min. After hydration, Zensis Axio Imager.Z2 was used for DIC to capture.

### Yeast Two-Hybrid

To test direct protein-protein interactions by Y2H, pairwise pGADT7 and pGBKT7 were co-transformed into yeast strain Y2H Gold (Clontech). Cells transformed with both plasmids were selected after growth 2 d at 30°C on synthetic dropout medium lacking leucine and tryptophan. Protein-protein interactions were then identified by growing the yeast for 2 d at 30°C on synthetic dropout medium lacking adenine, leucine, tryptophan, and histidine. To confirm interactions, single colony was diluted in sterile H_2_O to an OD600 of 0.1, and 10 μL was spotted onto both types of selective medium and again grown for 2 d at 30°C.

### Bimolecular Fluorescence Complementation (BiFC)

In planta protein-protein interactions were assayed by bimolecular fluorescence complementation in *N. benthamiana* leaves. The plasmids (pSYNE, YN; pSYCE, YC) were introduced into *Agrobacterium tumefaciens* strain *GV3101*. Overnight cultures were resuspended in 5 mL MMA buffer (0.15 M acetosyringone dilute in DMSO; 0.01 M MES, pH 7.5; 0.01 M MgCl_2_), incubated at room temperature for 4 h in the dark, and used for direct infiltration of 4 to 6 -week-old *N.benthamiana* leaves. Leaf sections of approximately 2×2 mm excised 36–48 h after infiltration were visualized under a Leica Stellaris 5 confocal laser scanning microscope using 40× objectives.

### Coimmunoprecipitation (Co-IP)

Co-IP in *N. benthamiana* was performed as described, with minor modifications (62). The *35S::ERf_CD_-GFP* and 3*5S::AP2σ-RFP* constructs were infiltrated into *N. benthamiana* via the *Agrobacterium*-mediated method. After 48 h, total proteins were extracted from the samples with IP buffer (50 mM Tris-HCl, pH 7.5; 150 mM NaCl; 5 mM dithiothreitol; 1% Triton X-100; 2% NP40 and 1×cocktail), followed by incubation with anti-GFP magnetic beads (D153-11, MBL) for 2 h. The beads were washed 6–8 times with washing buffer (50 mM Tris-HCl, pH 7.5; 150 mM NaCl; 1% Triton X-100; 2% NP40). The immunoprecipitated proteins were analyzed by SDS-PAGE and immunoblotted with anti-RFP antibody.

### Recombinant Protein Expression and Pull-down

To express recombinant proteins in prokaryotic cells, the coding sequences of the ERECTA_CD_, ERL1_CD_ and ERL2_CD_ were cloned into the pGEX4T-3 (GST tag) vector and transformed into *E.coli* strain *Rosetta*. GST-tagged proteins were purified with Glutathione Sepharose 4B beads (10250335, GE Healthcare) following the manufacturer’s instructions. AP2σ-MBP were purified using PurKine™ MBP-Tag Dextrin Resin 6FF (BMR20206, Abbkine). Protein concentration was determined using a BCA protein assay kit (Solarbio, PC0021).

The *in vitro* pull-down assay was performed as described in (62). MBP beads containing AP2σ-MBP and YFP-MBP were incubated with purified ERECTA_CD_-GST, ERL1_CD_-GST or ERL2_CD_-GST protein at 4°C for 2 h with gentle shaking. After washing several times with wash buffer, the proteins were detected by immunoblotting with anti-GST antibody.

### Pharmacological Treatment

For BFA (Sigma; B5936) treatment, Arabidopsis seedlings expressing GFP or YFP reporters were grown on 1/2 MS medium for 5 d, transferred to liquid medium with DMSO or 50 μM BFA for 1 or 2h and cell imaging of Arabidopsis seedlings.

For Wortmannin (Macklin; W820520-1mg) treatment, 5-day-old seedlings grown on 1/2 MS agar medium transferred to 1/2 MS liquid medium with DMSO or 50 μM Wm for 6h.

For ES9-17 (TargetMol; T64368) treatment was done as described in (63). Briefly, ES9-17 was dissolved as 50 mM stock using DMSO. The seedlings were grown on 1/2 MS containing 0.4% DMSO or 100 μM ES9-17 for 5 days and confocal imaged.

For Rhodamine B (Sigma; R6626) hexyl ester treatment, the protocol was performed as described (24). Briefly, 5-day-old seedlings were immersed into either mock (1% of DMSO) or 160 mM Rhodamine B hexyl ester solution for 30 min before imaging.

### Confocal Laser Scanning Microscopy

Confocal microscopy images were taken on the Leica Stellaris 5 inverted confocal microscope (Solms, Germany). For internalization imaging of ERECTA-YFP, ERL1-YFP and all other membrane organelle markers, maximum intensity projection of Z-stack images (0.33 μm step) covering the entire meristemoids were generated and subjected to analysis. The imaging was done with a 63×/1.2 W Corr lens on Leica Stellaris 5. A 514 nm laser was used to excite YFP and emission window of 518–600 nm was used to collect YFP signal. For the multicolor images of YFP and RFP, transient expression in tobacco leaves for 2 days were observed with a 40×/1.2 W Corr lens on Nikon A1R+ Ti2-E. 514 nm laser was used to excite YFP and 555 nm laser was used to excite RFP and FM4-64. Each experiment was repeated at least three times, each with multiple seedlings. Leica Stellaris 5 was used for assessing the effects of Wm and BFA on membrane organelle. For these purposes, images were taken using 100× glycerol immersion objective lens and 488/500–530 nm, 514/518–550 nm, 561/600–650 nm for GFP, YFP, and RFP signals, respectively with Time Gating of 0.3–0.6 ns to eliminate chloroplast autofluorescence. The Leica LAS AF software (http://www.leica-microsystems.com), and Fiji (https://imagej.net/Fiji) were used for post-acquisition image processing.

### MPK3/6 Phosphorylation Activity

For protein extraction, Arabidopsis samples were ground and homogenized in ice-cold extraction buffer (10 mM HEPES, pH 7.5; 100 mM NaCl; 1 mM EDTA pH 8.0; 10% Glycerol; 0.5% Triton X-100; 1 × cocktail). Samples were incubated on ice for 10–15 min and centrifuged at 4°C for 10 min at 12,000g. The supernatant was used for electrophoresis. For immune blotting, 20 mg of total proteins were separated by SDS-PAGE on 10% SDS-polyacrylamide gels, and MAPK activity was determined by the immune blot with phospho-P44/42 MAPK antibody (Cell Signaling Technology).

### Quantification and Statistical Analysis

All quantitative data were analyzed using either GraphPad Prism 10 or Microsoft Excel software. For comparisons between two experimental groups, two-tailed Student’s t-tests were employed. Multiple group comparisons were performed using one-way ANOVA followed by Tukey’s post hoc test. Statistical significance was defined as *p*< 0.05 for all analyses

## Supplemental Data Set

**Fig. S1.**
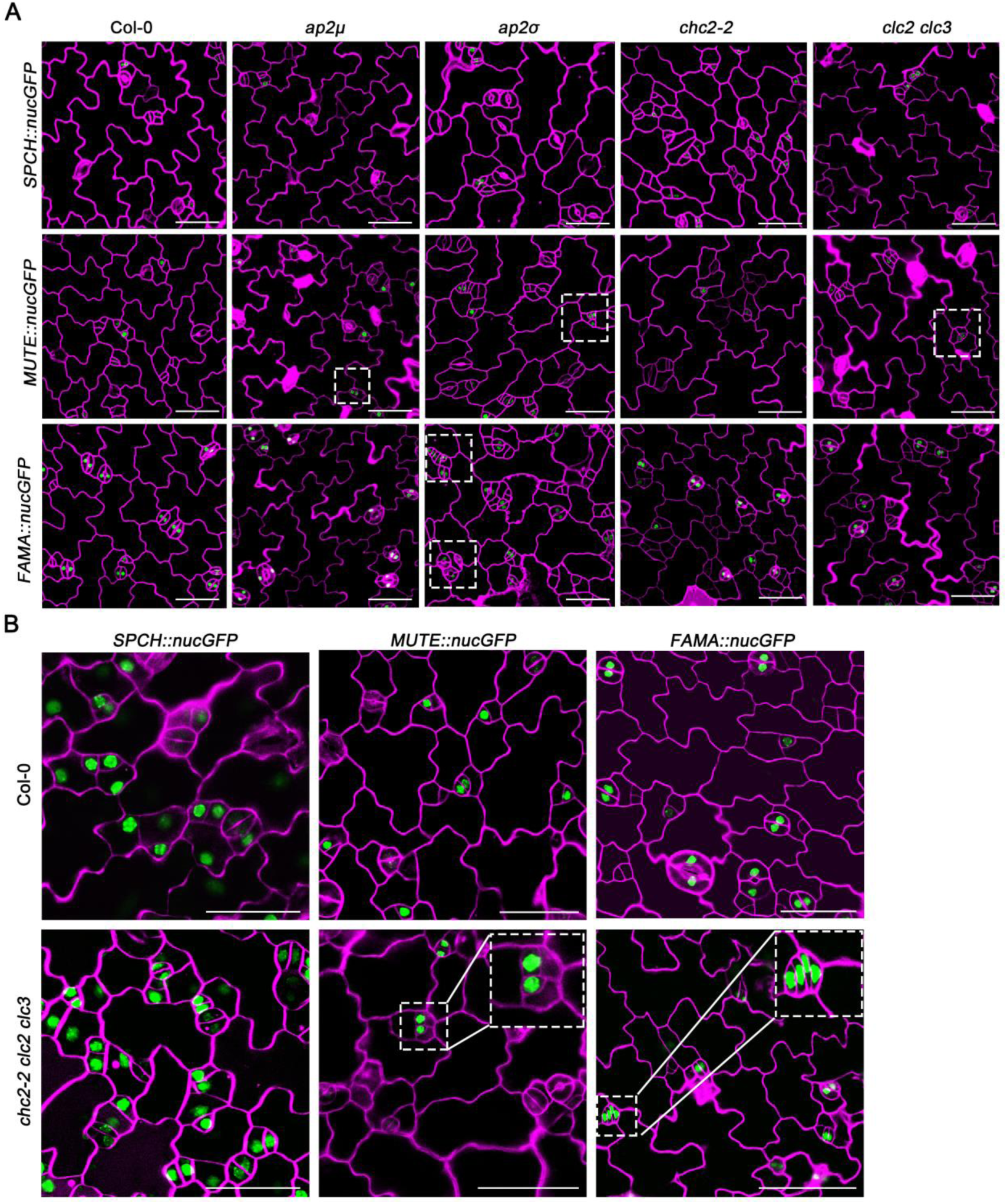
*ap2μ*, *ap2σ*, *chc2-2, clc2 clc3* and *chc2-2 clc2 clc3* mutants showed disordered stomatal development patterning. (A) Expression patterning of *pSPCH::nucGFP*, *pMUTE::nucGFP* and *pFAMA::nucGFP* in WT, *ap2μ*, *ap2σ*, *chc2-2* and *clc2 clc3* seedlings grown on 1/2 MS medium for 5 days. Red fluorescence is FM4-64 staining, and white dotted boxes indicate abnormally developed cells. Scale bars, 50 μm. (B) Expression patterning of *pSPCH::nucGFP*, *pMUTE::nucGFP* and *pFAMA::nucGFP* in WT and *chc2-2 clc2 clc3* grown on 1/2 MS medium for 5 days. Magenta is FM4-64 staining, and the white dotted box indicates abnormally developed cells. Scale bars, 50 μm.

**Fig. S2.**
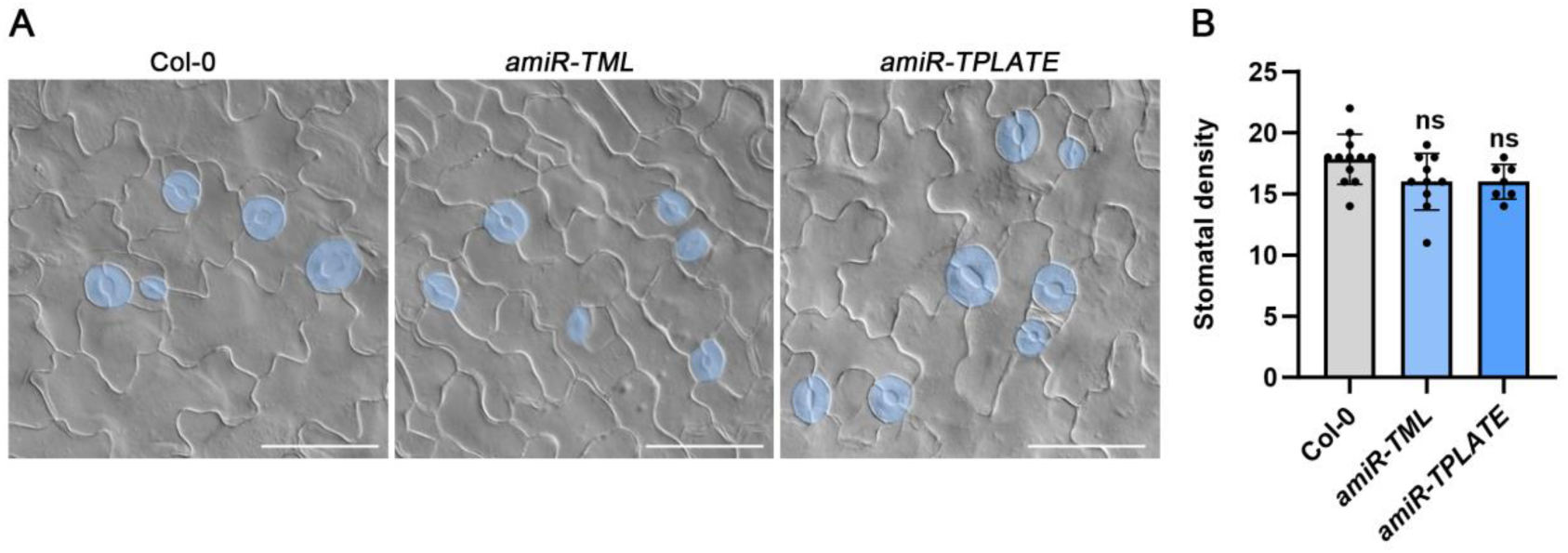
TPC did not involve in CME-mediated stomatal development. (A) DIC images showed abaxial epidermal cell patterns on cotyledons at 5 dpg in Col-0, *amiR-TML* and *amiR-TPLATE* induced by 20 μM ESTR. For easy identification, the stomata are painted blue. Scale bars, 50 μm. (B) Statistical chart showed the number of stomata of Col-0, *amiR-TPLATE* and *amiR-TML* at 0.1 mm^2^ area. Error line represents the SE. *T*-test is used for significance analysis, ns represents *P*>0.05, no significant difference.

**Fig. S3.**
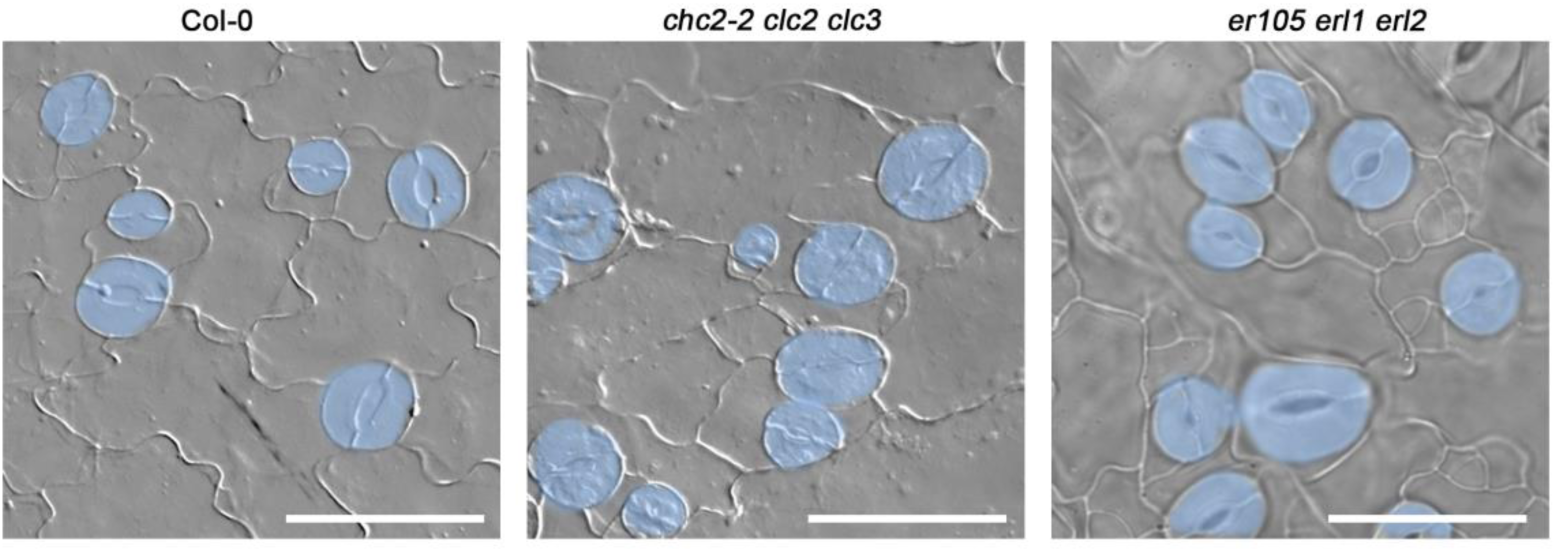
The stomatal clustering phenotype of *chc2-2 clc2 clc3* resembled that of *er105 erl1 erl2* mutants. DIC images showed abaxial epidermal cell patterns on cotyledons at 5 dpg in Col-0, *chc2-2 clc2 clc3* and *er105 erl1 erl2.* For easy identification, the stomata are painted blue. Scale bars, 50 μm.

**Fig. S4.**
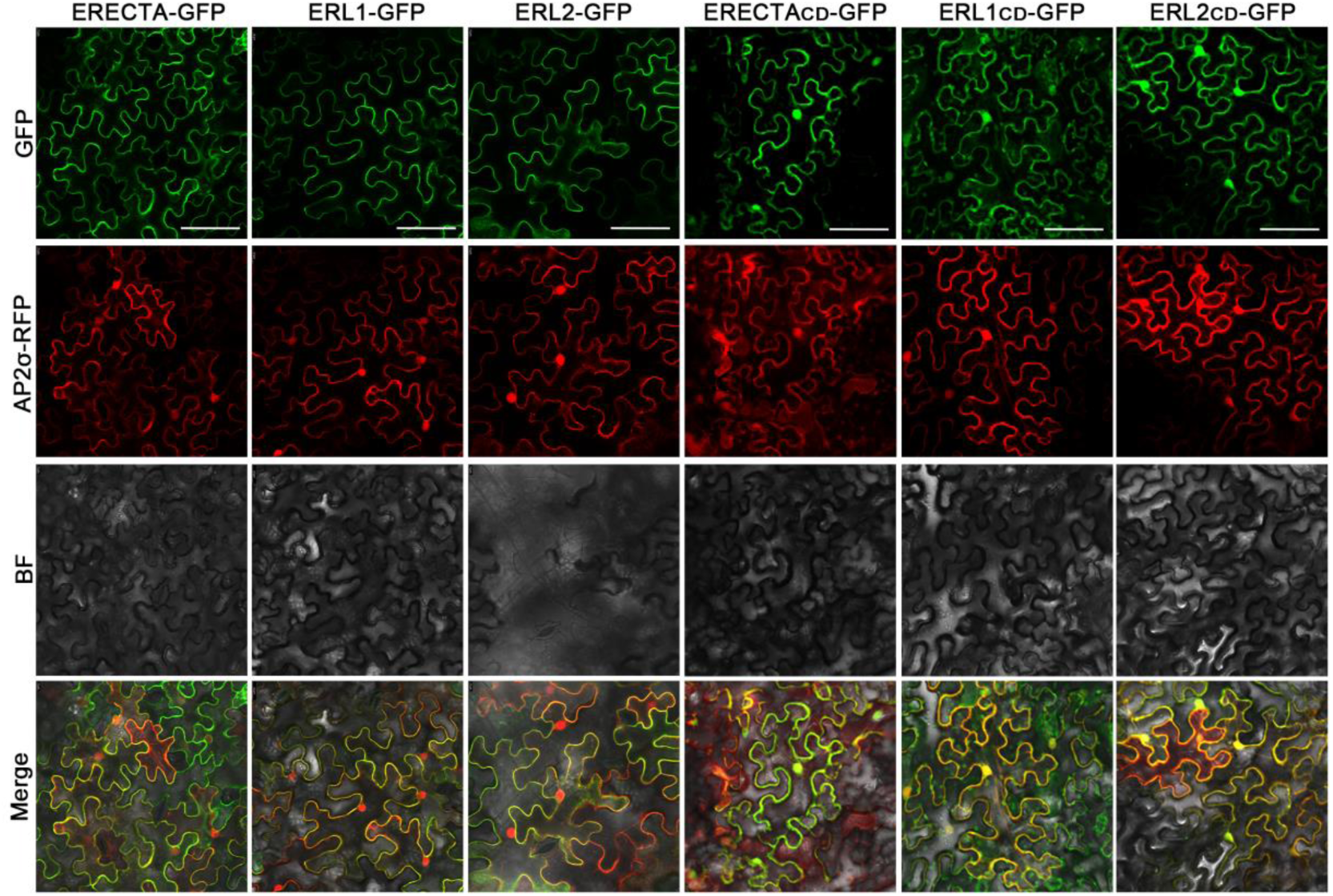
ERf and AP2σ were co-localized in the plasma membrane, and ERf_CD_ and AP2σ were co-localized in the plasma membrane and nucleus. Confocal images showed that the expression patterning of ERECTA, ERL1, ERL2 and AP2σ in tobacco epidermal cells. ERf and ERf_CD_ were marked GFP, AP2σ was marked RFP, BF represents bright field, merge is fusion of GFP and RFP. Scale bars, 50 μm.

**Fig. S5.**
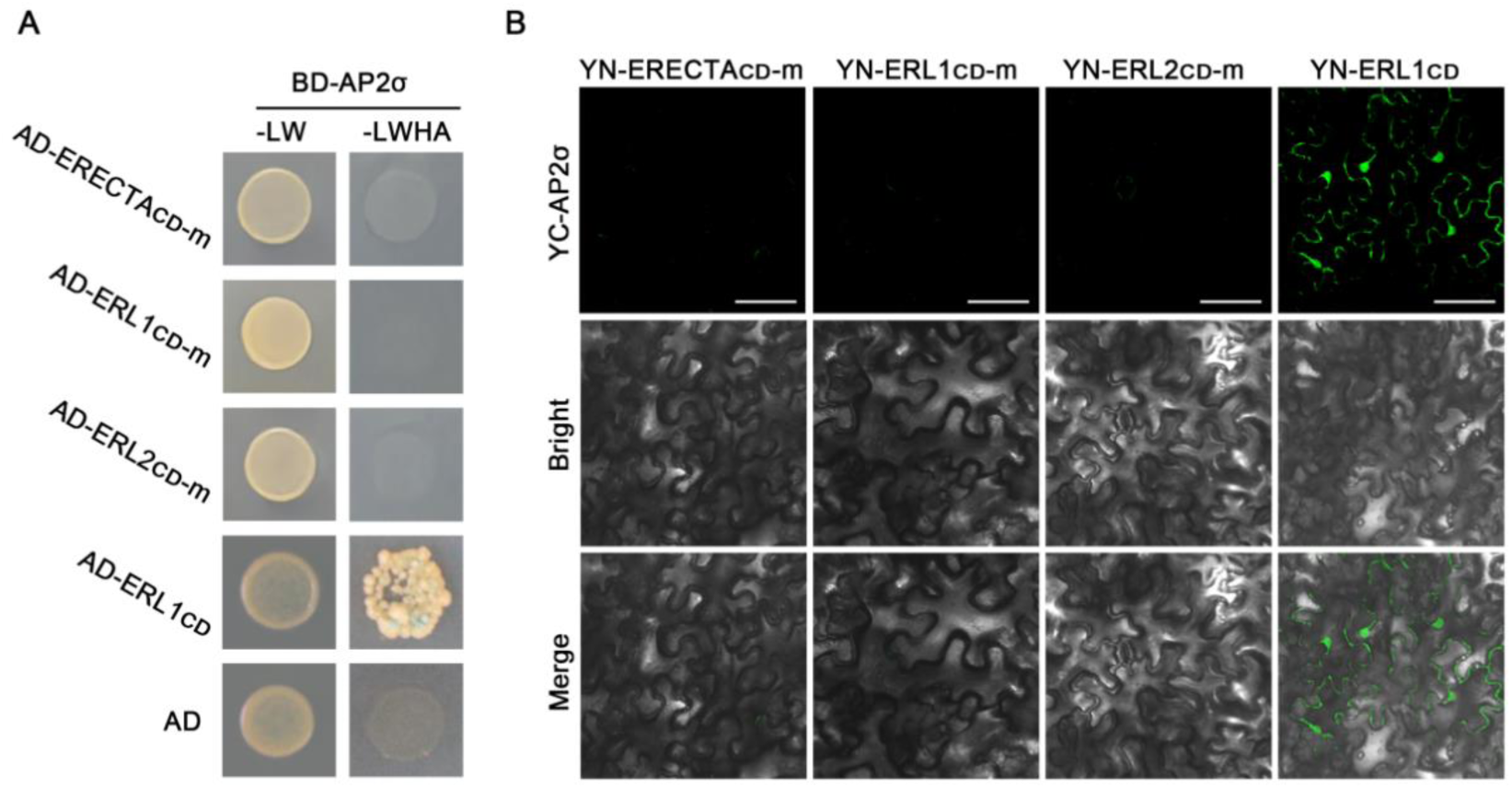
AP2μ subunit can’t interacted with ERECTA_CD_, ERL1_CD_ and ERL2_CD_. (A) Y2H assayed the interaction of AP2μ with ERECTA_CD_-m, ERL1_CD_-m and ERL2_CD_-m. “-LW” represents yeast two-deficiency medium (SD/-Trp/-Leu), “-LWHA” represents four-deficiency medium (SD/-Trp/-Leu/-His/-Ade), ERL1_CD_ was used as a positive control and AD represents pGADT7 empty. (B) BiFC verified the interaction of AP2 μ with ERECTA_CD_-m, ERL1_CD_-m and ERL2_CD_-m, ERL1_CD_ was used as a positive control. Scale bars, 100 μm.

**Fig. S6.**
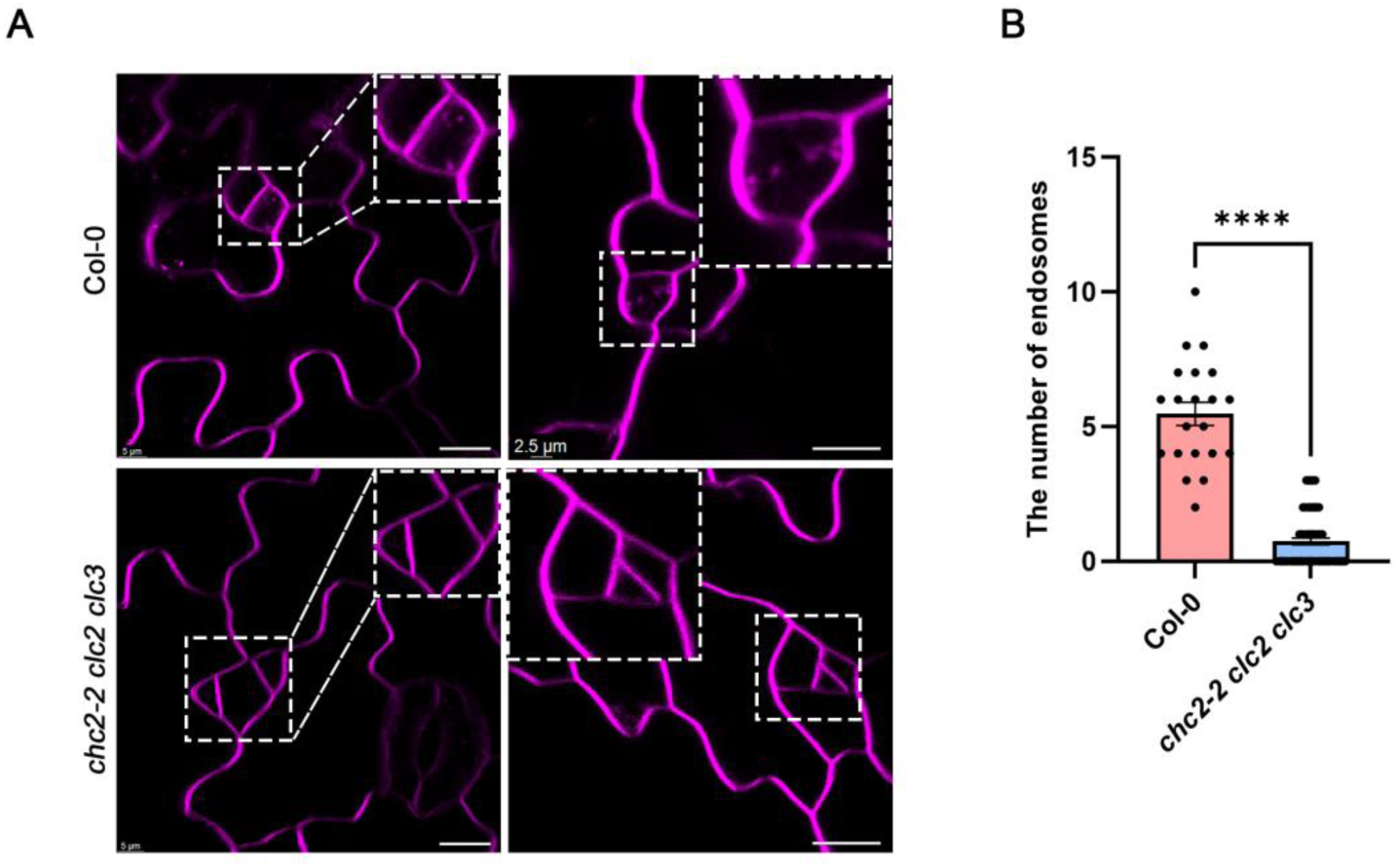
Endocytosis defected in cotyledon epidermal cells of *chc2-2 clc2 clc3* mutant. (A) Confocal images showed WT and *chc2-2 clc2 clc3* mutant cotyledon epidermal cells stained with 2 μM FM4-64 for 5 days and imaged immediately after 30 min. Scale bars, 10 μm. (B) Quantitative analysis of the number of endocytic bodies in WT and *chc2-2 clc2 clc3* mutants. Thirty cells from 5 cotyledons were measured. The error line represents the SE. *T*-test is performed significance analysis. **** indicates *P*<0.0001.

**Fig. S7.**
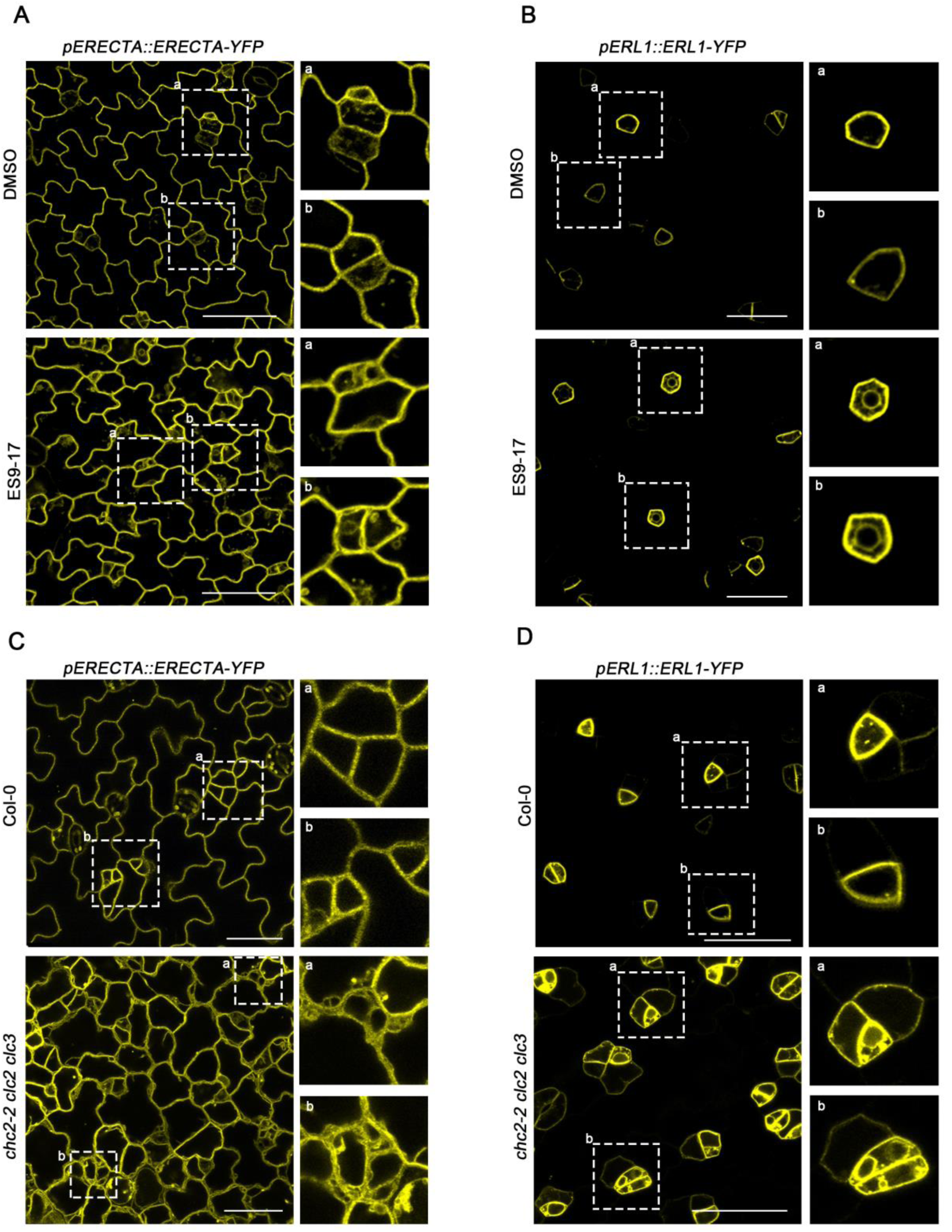
Both CME inhibitor ES9-17 treatment and clathrin deficient mutant changed the subcellular localization of ERECTA and ERL1. (A-B) Confocal images showed that *pERECTA::ERECTA-YFP* and *pERL1::ERL1-YFP* lines were treated with 50 μM ES9-17 for 5 d. After ES9-17 treatment, ERECTA and ERL1 generate ring-like structures in the cytoplasm. DMSO was used as the control, with an addition of 0.1%. The right figure is an enlarged view of the white dashed box in the left figure. (C-D) Confocal images showed that the subcellular localization of ERECTA and ERL1 in *chc2-2 clc2 clc3* mutants. The picture on the right side is an enlarged picture of the white dotted box. Scale bars, 20 μm

**Fig. S8.**
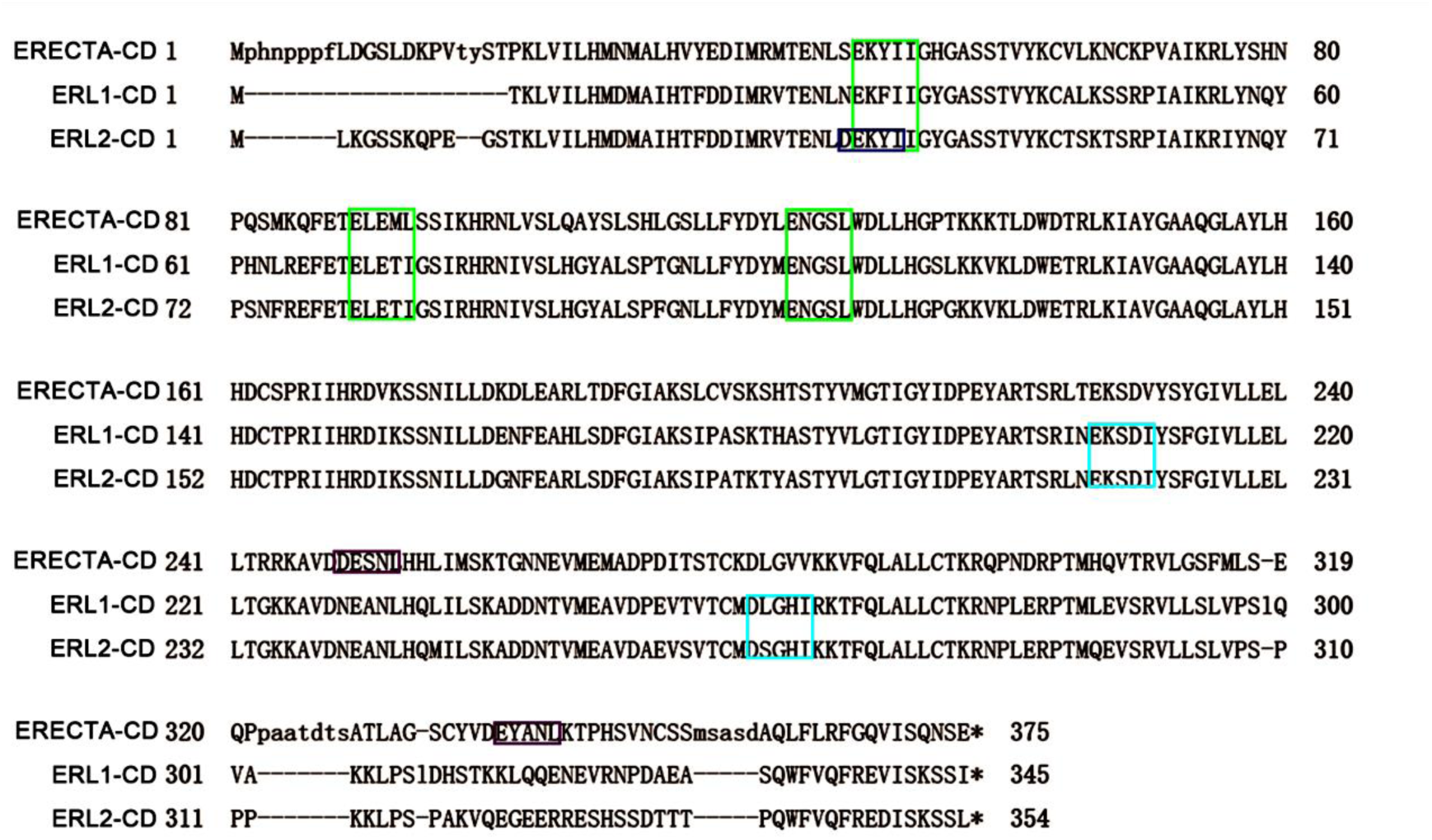
Analysis of endocytosis motifs in the intracellular region of ERECTA, ERL1 and ERL2. The green box represents ERECTA, ERL1 and ERL2; the blue box represents an endocytic motif specific to ERL2; the cyan box represents an endocytic motif unique to ERL1 and ERL2; and the purple box represents an endocytic motif unique to ERECTA.

**Fig. S9.**
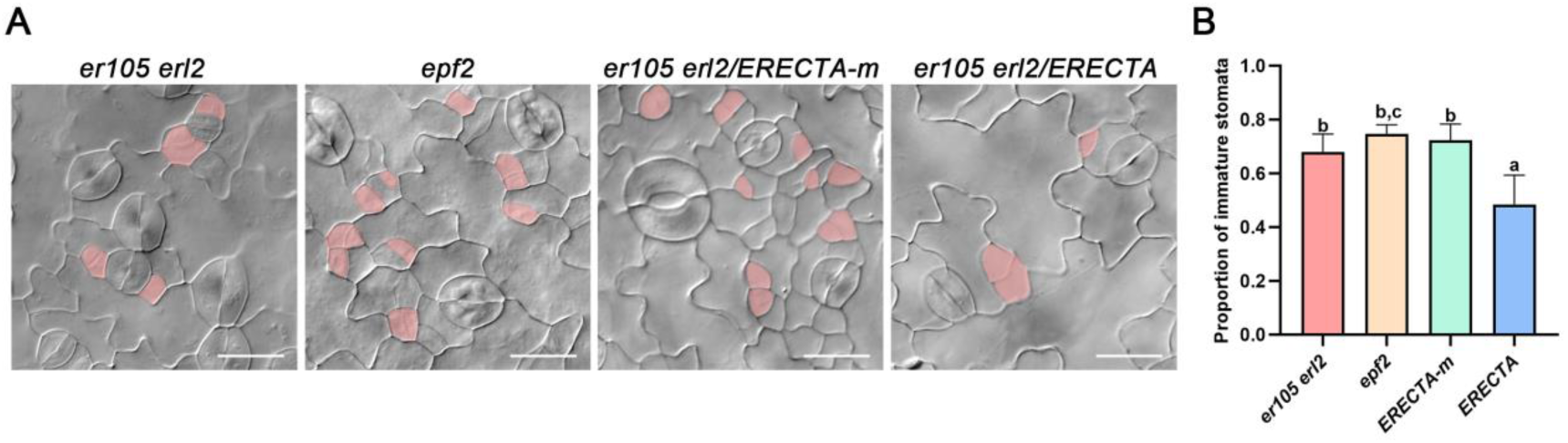
The mutation in the endocytosis motif of ERECTA fails to rescue the defects of *er105 erl2*. (A) DIC image showed that adaxial epidermal cell patterns on cotyledons at 5 dpg in *er105 erl2* (top left column), *epf2* (top middle column), *ERECTA-m-YFP/er105 erl2* (top right column), *ERECTA-YFP/er105 erl2* (bottom left column). *ERECTA-m* cannot complement the phenotype of *er105 erl2* and exhibits a phenotype similar to that of *epf2*. For easy observation, the MMC or M is painted blue. Scale bars, 50 μm. (B) Quantitative analysis showed that the proportion of immature stomata in *er105 erl2*, *epf2*, *ERECTA-m/er105 erl2*, and *ERECTA/er105 erl2*. Tukey test is determined for significance, values are mean ± SEM (n > 5).

**Tab. S1.**
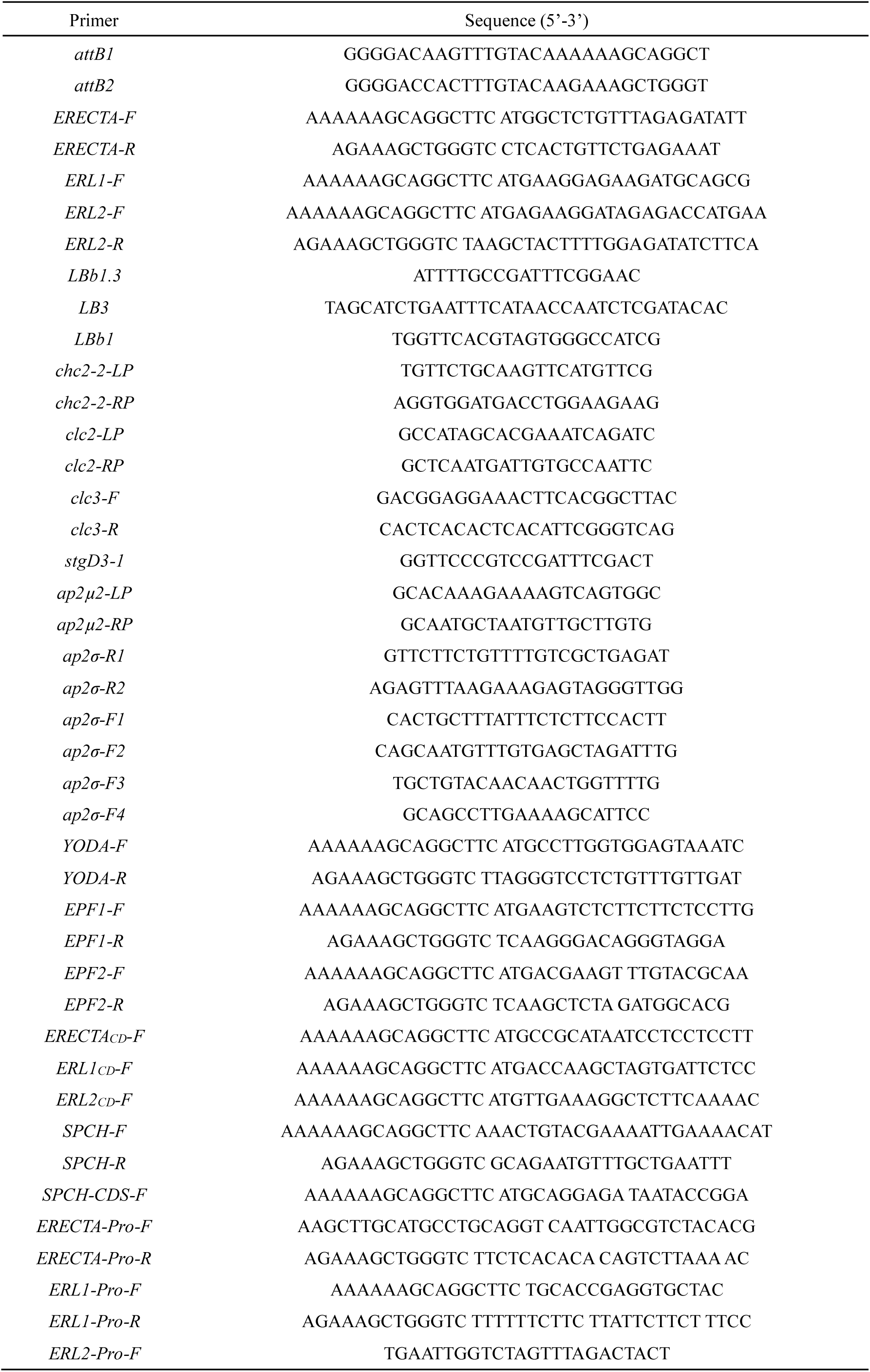
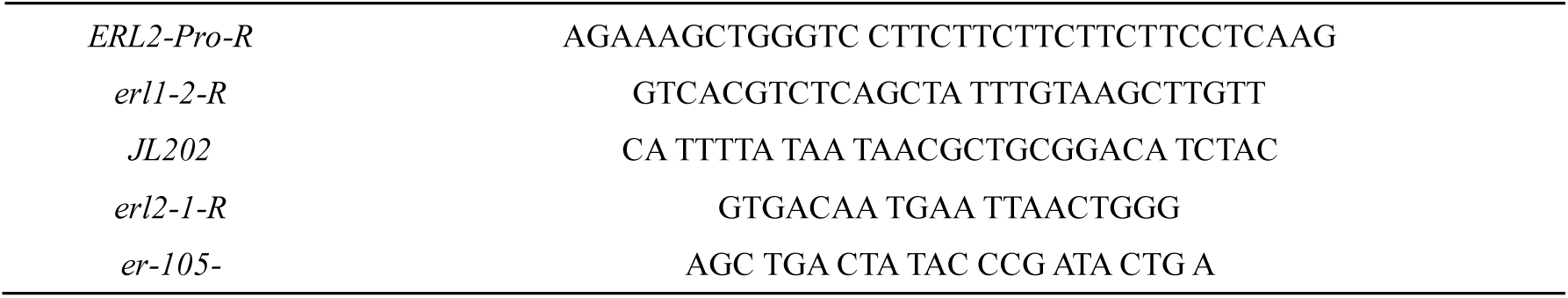
Primer Sequence.

